# Loss of TMEM106B exacerbates Tau pathology and neurodegeneration in PS19 mice

**DOI:** 10.1101/2023.11.11.566707

**Authors:** Tuancheng Feng, Huan Du, Fenghua Hu

## Abstract

*TMEM106B*, a gene encoding a lysosome membrane protein, is tightly associated with brain aging, hypomyelinating leukodystrophy, and multiple neurodegenerative diseases, including frontotemporal lobar degeneration with TDP-43 aggregates (FTLD-TDP). Recently, *TMEM106B* polymorphisms have been associated with tauopathy in chronic traumatic encephalopathy (CTE) and FTLD-TDP patients. However, how TMEM106B influences Tau pathology and its associated neurodegeneration, is unclear. Here we show that loss of TMEM106B enhances the accumulation of pathological Tau, especially in the neuronal soma in the hippocampus, resulting in severe neuronal loss in the PS19 Tau transgenic mice. Moreover, *Tmem106b^-/-^* PS19 mice develop significantly increased disruption of the neuronal cytoskeleton, autophagy-lysosomal function, and lysosomal trafficking along the axon as well as enhanced gliosis compared with PS19 and *Tmem106b^-/-^* mice. Together, our findings demonstrate that loss of TMEM106B drastically exacerbates Tau pathology and its associated disease phenotypes, and provide new insights into the roles of TMEM106B in neurodegenerative diseases.

## Introduction

*TMEM106B* was first identified as a genetic risk factor for frontotemporal lobar degeneration (FTLD) with TDP-43 aggregates [20,30,88,89]. Subsequently, *TMEM106B* polymorphisms (SNP rs1990622) have been associated with FTLD caused by *C9ORF72* mutations [21,32,47,87], cognitive impairment in amyotrophic lateral sclerosis (ALS)[90], Alzheimer’s disease (AD) [77], Parkinson’s disease (PD) [86], chronic traumatic encephalopathy (CTE) [8,17], hippocampal sclerosis in aging (HS-Aging) [64,97], limbic-predominant age-related TDP-43 encephalopathy (LATE) [63], and brain aging [76]. Moreover, a dominant D252N mutation in TMEM106B causes hypomyelinating leukodystrophy (HLD) [83,95]. In addition, the TMEM106B rs1990622 risk allele is also associated with increased Purkinje cell degeneration in humans [27]. Intriguingly, C-terminal fragments of TMEM106B were recently shown to form amyloid fibrils in the aged brain and the brains of patients with a diverse set of neurodegenerative diseases [14,19,25,42,69,70,79].

A recent study has found an association between the TMEM106B risk allele (SNP rs3173615 T185) and increased phosphorylated Tau (AT8) density, neuroinflammation, synaptic dysfunction, and ante-mortem dementia in persons with CTE [17]. Additionally, TMEM106B rs1990622 A/A risk allele is linked to distinctive medial temporal predominant, 4-repeat, neuro-astroglial tauopathy in FTLD/ALS-TDP patients [52]. These studies suggest that TMEM106B might influence Tau pathology and Tau-mediated neurodegeneration.

TMEM106B is a type II transmembrane protein localized in late endosome/lysosome compartments [11,15,46]. A large body of evidence demonstrates that TMEM106B is critical for proper lysosomal functions, via influencing lysosomal morphology [11,15,46,84], lysosome pH [15,43,45], lysosomal enzyme activity [43,53], lysosome exocytosis [45], lysosomal positioning [29], lysosomal trafficking in dendrites [80] and axons [27,28,53].

Tau is a microtubule-associated protein that stabilizes microtubules (MT) and promotes microtubule assembly [55]. Tauopathies characterized by cytoplasmic aggregation of hyperphosphorylated, and filamentous Tau protein are seen in several neurodegenerative diseases in humans, including Alzheimer’s disease, FTLD, and CTE [75]. Mutations in the Tau-encoding gene *MAPT* cause approximately 5% of FTLD cases [34]. Hyperphosphorylated and aggregated Tau due to mutation in the *MAPT* gene leads to microtubule disruption, axonal transport defects, neuroinflammation, neuronal loss, and cognitive deficits [1–3,6,12,16,18,78].

To determine the effect of TMEM106B on Tauopathy, we ablated TMEM106B in the PS19 tauopathy mouse model, in which P301S human Tau (hTau) protein containing 4 carboxyl-terminal repeats (4R) is expressed under the prion promoter [96]. Our data demonstrate that TMEM106B deficiency in the PS19 mice causes a dramatic increase of Tau and phosphorylated Tau in the sarkosyl-insoluble fraction and alternations in the subcellular distribution of Tau. In addition, these mice exhibit severe neuron loss, glia activation, and brain atrophy, as well as exacerbated lysosomal-autophagy abnormalities. Taken together, these results provide novel insights into the role of TMEM106B in Tauopathy, brain aging, and neurodegenerative diseases.

## Results

### TMEM106B depletion exacerbates neurodegeneration and motor dysfunction in PS19 mice

To determine whether TMEM106B affects Tau pathology and Tauopathy-mediated neurodegeneration, we crossed *Tmem106b^-/-^* mice [29] with the PS19 mice [96] to generate *Tmem106b^-/-^* PS19 mice. To limit variabilities in phenotypes in male and female PS19 mice [85], only male mice were used in our studies. Strikingly, we observed significant brain atrophy in 8.5-month-old *Tmem106b^-/-^* PS19 mice, but not in the brains of PS19, *Tmem106b^-/-^*, and WT controls (Fig. 1A). A drastic decline in the hippocampal volume was detected in 8.5-month-old *Tmem106b^-/-^* PS19 mice (Fig. 1B and 1C), which is highly correlated with neuronal loss observed in the pyramidal cell layer in the hippocampus CA3 region of 8.5-month-old *Tmem106b^-/-^* PS19 mice (Fig. 1D and 1E). Consistent with the immunostaining result, western blot analysis revealed a significant decrease in the levels of neuronal marker NeuN in the whole brain lysate of 8.5-month-old *Tmem106b^-/-^* PS19 mice (Supplementary Fig. 1A and 1B). To further examine the cause of neuronal death, we stained brain sections with antibodies against cleaved caspase-3 (c-caspase-3), a key mediator of apoptosis, and detected a significant increase of c-caspase-3 in the hippocampus, especially the fornix, the white matter tract that connects the hippocampus to several subcortical brain regions [5], in 8.5-month-old *Tmem106b^-/-^* PS19 mice, compared to the PS19 mice (Supplementary Fig. 1D and 1E). Moreover, immunostaining with antibodies against axon marker NF-L showed a significant loss of axons in the hippocampus of 8.5-month-old *Tmem106b^-/-^* PS19 mice, compared to the PS19 mice (Fig. 1F and 1G), suggesting axonal degeneration in the brain of these mice.

**Figure 1.**
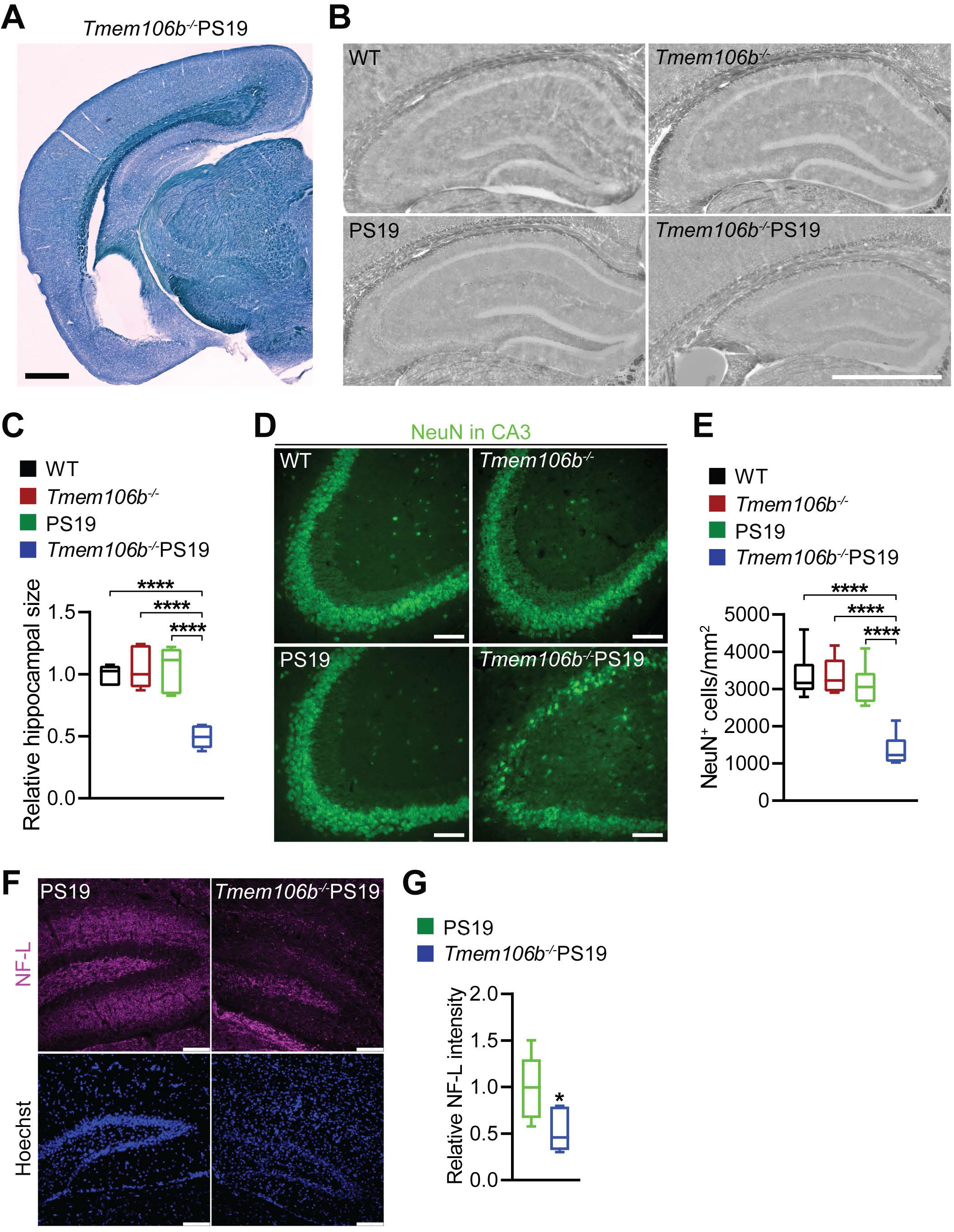
TMEM106B depletion exacerbates neurodegeneration in PS19 mice. **(A)** Representative image of 8.5-month-old*Tmem106b^-/-^* PS19 mice brain section stained with Trueblack. Scale bar: 0.5 mm. **(B, C)** Representative bright-field images of the hippocampus in 8.5-month-old WT, *Tmem106b^-/-^*, PS19, and *Tmem106b^-/-^* PS19 mice. The area of the hippocampus was quantified in C. n=6. Scale bar: 1 mm. Data presented as mean ± SEM. One-way ANOVA tests with Bonferroni’s multiple comparisons: ****, p<0.0001. **(D, E)** Representative images of NeuN immunostaining in the hippocampal CA3 region in 8.5-month-old WT, *Tmem106b^-/-^*, PS19, and *Tmem106b^-/-^* PS19 mice. The density of NeuN-positive neurons was quantified in E. n=6. Scale bar: 100 µm. Data presented as mean ± SEM. One-way ANOVA tests with Bonferroni’s multiple comparisons: ****, p<0.0001. Scale bar: 100 µm. **(F, G)** Representative images of NF-L immunostaining in the hippocampal dentate gyrus (DG) region in 8.5-month-old WT, *Tmem106b^-/-^*, PS19, and *Tmem106b^-/-^* PS19 mice. The intensity of NF-L was quantified in G. n=6. Scale bar: 100 µm. Data presented as mean ± SEM. Unpaired Student’s t-test: *, p<0.05.

### TMEM106B ablation exaggerates Tau pathology in PS19 mice

To examine the effect of TMEM106B deletion on Tau pathology, we sequentially extracted Tau from the mouse brain using the high salt buffer and 1% sarkosyl buffer and analyzed Tau levels in different fractions by western blot. At 8.5 months of age, PS19 mice are known to accumulate aggregated Tau. Compared to PS19 mice, *Tmem106b^-/-^* PS19 mice have significantly higher levels of hTau and phosphorylated Tau (Ser-404 and Thr-205) in the sarkosyl-insoluble fraction (Fig. 2A and 2B), whereas no obvious changes in the level of soluble Tau were detected in sarkosyl-soluble fraction (Fig. 2A and 2B). Moreover, the increase in the levels of hTau and phosphorylated Tau (Ser-404) was observed in the sarkosyl-insoluble fraction from the hippocampus of 5 to 5.4-month-old young *Tmem106b^-/-^*PS19 mice (Supplementary Fig. 2A and 2B). Consistently, a dramatic increase of hTau and phosphorylated Tau (Ser-404) intensities was detected in the hippocampus of 8.5-month-old *Tmem106b^-/-^* PS19 mice, compared to PS19 mice by immunofluorescence staining (Fig. 2C and 2D).

**Figure 2.**
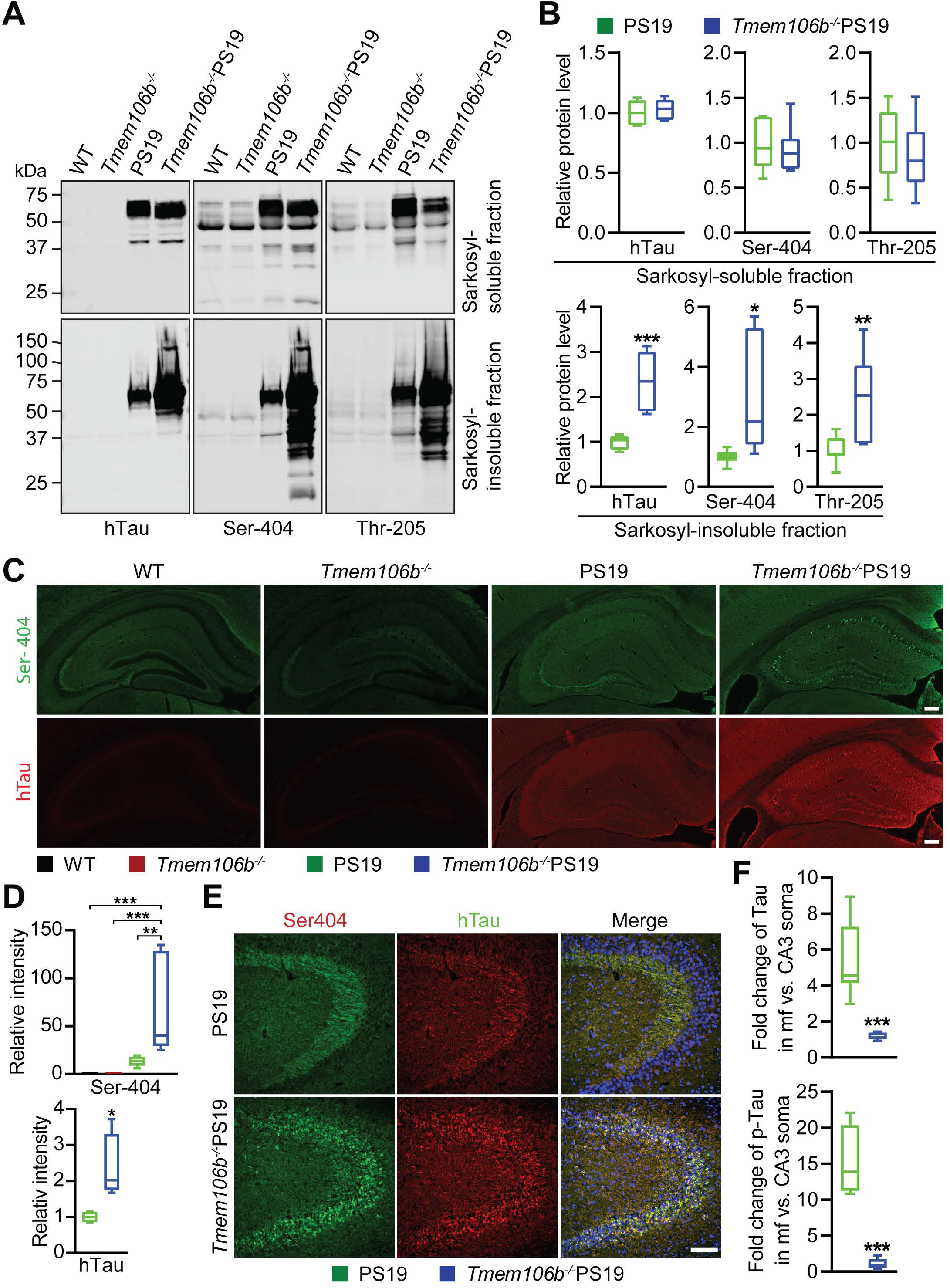
TMEM106B depletion accelerates Tau pathology and leads to neuronal somatic accumulation of hTau and phosphorylated Tau in PS19 mice. **(A, B)** Western blot analysis of hTau and phosphorylated Tau (Ser-404) and Thr-205 in sarkosyl-soluble fraction and sarkosyl-insoluble fraction extracted from the brain of 8.5-month-old WT, *Tmem106b^-/-^*, PS19, and *Tmem106b^-/-^* PS19 mice. Relative levels of indicated proteins were quantified in B. n=5-7. Data presented as mean ± SEM. Unpaired Student’s t-test: *, p<0.05; **, p<0.01; ***, p<0.001. **(C, D)** Representative images of phosphorylated Tau (Ser-404) or hTau immunostainings in the hippocampus of 8.5-month-old WT, *Tmem106b^-/-^*, PS19, and *Tmem106b^-/-^* PS19 mice brain sections. Relative Ser-404 and hTau intensity in the hippocampus were quantified in D. n=4-6. Scale bar: 100 µm. Data presented as mean ± SEM. One-way ANOVA tests with Bonferroni’s multiple comparisons: *, p<0.05; **, p<0.01; ***, p<0.001. **(E, F)** Representative images of phosphorylated hTau (Ser-404) and hTau co-immunostainings in the hippocampus of 8.5-month-old WT, *Tmem106b^-/-^*, PS19, and *Tmem106b^-/-^* PS19 mice. Fold change of hTau or Ser-404 intensity in mossy fiber vs. CA3 neuronal soma were quantified in F. n=5-6. Scale bar: 100 µm. Data presented as mean ± SEM. Unpaired Student’s t-test: *, p<0.05.

Interestingly, while PS19 mice accumulate hTau and phosphorylated Tau (Ser-404) signals in the mossy fibers, the axons of dentate gyrus (DG) granule cells, *Tmem106b^-/-^* PS19 mice show intense Tau signals in the neuronal soma in the pyramidal cell layer in the CA1 and CA3 regions, and the granule cell layer in the DG (Fig. 2C, 2E and 2F), indicating a significant redistribution of pathological hTau and phosphorylated Tau (Ser-404) from axons to soma at 8.5 months of age when TMEM106B is ablated. However, 5 to 5.4-month-old *Tmem106b^-/-^* PS19 mice do not exhibit the altered distribution of Tau in the hippocampus (Supplementary Fig. 2C), suggesting that the redistribution of Tau is age-dependent and might be downstream of other events.

To determine whether TMEM106B affects the homeostasis of mouse Tau at endogenous levels, we performed western blot analyses of the lysates extracted from the hippocampus and brains of WT and *Tmem106b^-/-^* mice at different ages. Loss of TMEM106B does not affect the levels of endogenous mouse Tau and phosphorylated mouse Tau (Ser-404) in mice of 5-month-old or 16-month-old (Supplementary Fig. 3A-D). Together, these data suggested that loss of TMEM106B in mice exaggerates Tau pathology specifically in PS19 mice with overexpressed human 4R Tau.

### TMEM106B ablation exacerbates alterations in neuronal microtubule structure in the PS19 mice

The redistribution of hTau from axons to soma in the *Tmem106b^-/-^* PS19 mice (Fig. 2C, 2E and 2F) suggests that the trafficking of Tau might be affected by TMEM106B ablation. Microtubules (MT), comprised of α- and β-tubulin heterodimers, are one of the three main cytoskeleton components and serve as tracks for motor protein-based bidirectional transport of cargo [72,81]. Altered microtubule function in axons usually results in protein trafficking defects [6,13,38]. Tau pathology has been shown to affect microtubule dynamics and axonal transport due to the dissociation of Tau from microtubules [1]. PS19 mice have been reported to exhibit a progressive alteration of α-tubulin in the hippocampus starting at 3 months old [96]. To determine changes in MT structure in the *Tmem106b^-/-^* PS19 mice, immunostaining was performed using antibodies against β-III tubulin and hTau. We found an accumulation of abnormal β-III-tubulin punctas in neuronal soma in the hippocampal CA3 region of 8.5-month-old *Tmem106b^-/-^* PS19 mice (Fig. 3A).

**Figure 3.**
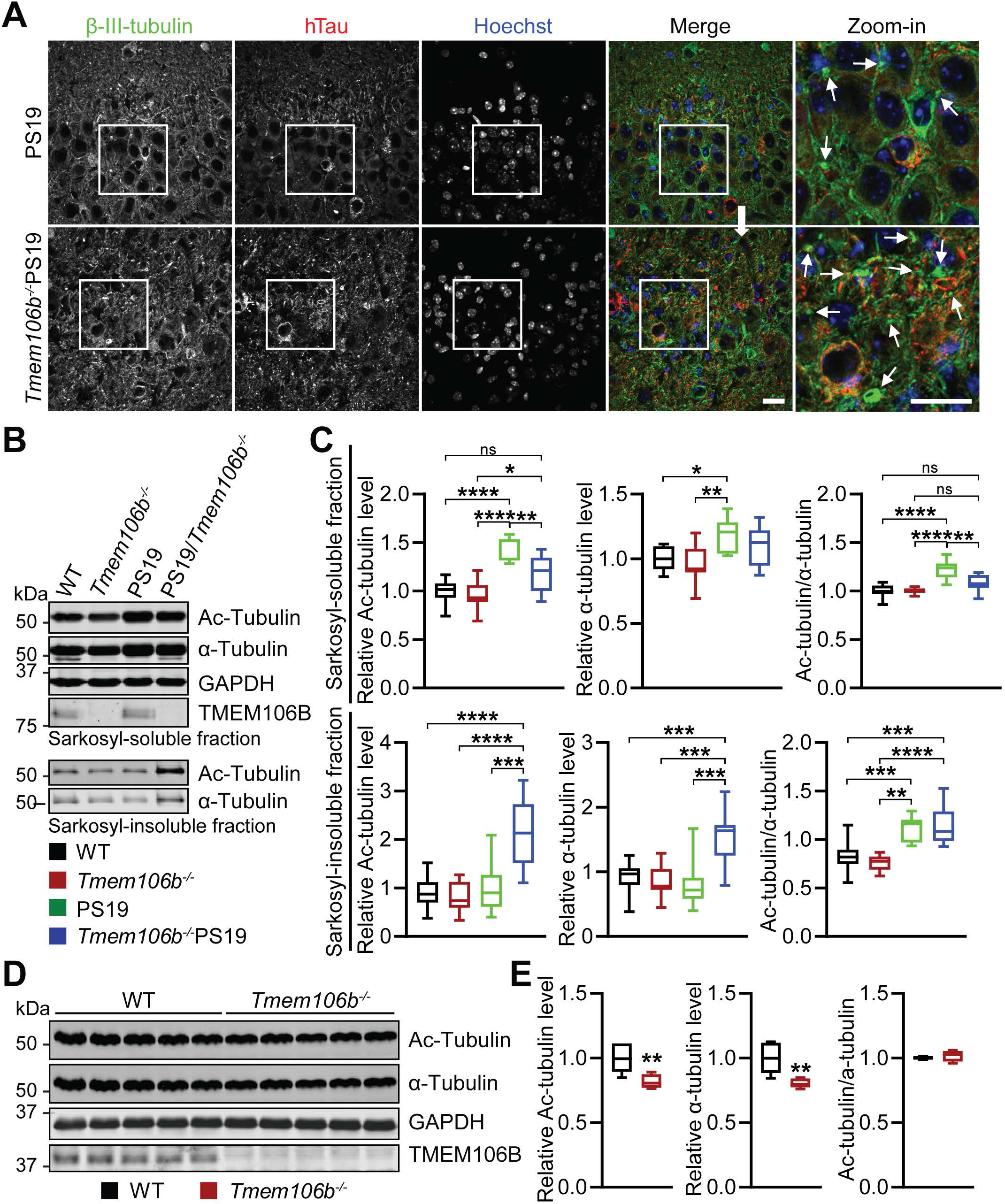
TMEM106B depletion exacerbates alterations in microtubule structure in neurons in PS19 mice. **(A)** Representative images of β-III-tubulin and hTau co-immunostainings in CA3 in the hippocampus of 8.5-month-old PS19 and *Tmem106b^-/-^* PS19 mice brain sections. Zoom-in images show the accumulation of β-III-tubulin punctas as indicated by white arrows in CA3 in the hippocampus of 8.5-month-old *Tmem106b^-/-^* PS19 mice. n=6. Scale bar: 20 µm. **(B, C)** Western blot analysis of Ac-tubulin, α-tubulin, TMEM106B, and GAPDH in sarkosyl-soluble fraction and sarkosyl-insoluble fraction extracted from the brain of 5 to 5.4-month-old WT, *Tmem106b^-/-^*, PS19, and *Tmem106b^-/-^* PS19 mice. Relative levels of indicated proteins were quantified in C. n=7-12. Data presented as mean ± SEM. One-way ANOVA tests with Bonferroni’s multiple comparisons: *, p<0.05; **, p<0.01; ***, p<0.001; ****, p<0.0001. n.s.: non-significant. **(D, E)** Western blot analysis of Ac-tubulin, α-tubulin, TMEM106B, and GAPDH in RIPA-soluble fraction from 16-month-old WT and *Tmem106b^-/-^* mice brain. Relative levels of indicated proteins were quantified in E. n=5. Data presented as mean ± SEM. Unpaired Student’s t-test: **, p<0.01.

Acetylation of Lys-40 on α-tubulin regulates MT dynamics and stability and is strongly associated with long-lived MTs [23,73]. A reduction of acetylated alpha-tubulin (Ac-tubulin) was found in neurofibrillary tangle-bearing neurons in Alzheimer’s disease [39]. Next, we examined the protein level of Ac-tubulin and α-tubulin by western blot and found that the protein levels of Ac-tubulin and the ratio of Ac-tubulin/α-tubulin are significantly lower in the sarkosyl-soluble fraction in the *Tmem106b^-/-^* PS19 mice when compared to PS19 mice, although both PS19 and *Tmem106b^-/-^* PS19 mice show increased levels of Ac-tubulin and α-tubulin compared to *Tmem106b^-/-^* and WT control (Fig. 3B and 3C). Interestingly, in sarkosyl-insoluble fraction, the protein levels of Ac-tubulin and α-tubulin are significantly increased in *Tmem106b^-/-^* PS19 mice, compared to PS19, *Tmem106b^-/-^* and WT controls (Fig. 3B and 3C). The changes in Ac-tubulin and α-tubulin levels and solubility in *Tmem106b^-/-^* PS19 mice compared to PS19 mice suggest a possible role of TMEM106B in regulating tubulin dynamics. However, we failed to detect a consistent change in tubulin in 5 to 5.4-month-old *Tmem106b^-/-^* mice due to variability (Fig. 3B and 3C). To further probe whether TMEM106B deficiency alone affects tubulin, we analyzed Ac-tubulin and α-tubulin levels in the brain lysates of 16-month-old *Tmem106b^-/-^* mice and found a modest but significant decrease in Ac-tubulin and α-tubulin levels in the soluble fraction when compared to age-matched WT controls (Fig. 5D. 5E). Taken together, these results support a role of TMEM106B in regulating tubulin dynamics.

### TMEM106B deletion leads to the accumulation of abnormal NF-containing aggregates in PS19 mice

Neurofilaments (NF), another cytoskeleton component in axons, are comprised of neurofilament light chain (NF-L), neurofilament medium chain (NF-M), neurofilament heavy chain (NF-H), alpha-internexin (INA), and peripherin (PRPH) [99]. A large amount of evidence demonstrated that NF-L, often used as a biomarker in neurodegenerative diseases, is involved in the pathophysiological processes underlying different states of neurodegeneration[10,31,98]. Overexpression of NF-L in mice results in neurofilament accumulation accompanied by axonal swelling and degeneration in motor neurons [94], whereas loss of NF-L in mice leads to the formation of NF-positive aggregates in motor neurons [50], and motor neuron death [58]. The phosphorylation of the NF-H tail domain plays a central role in modulating NF organization and regulates Tau trafficking along the axons [10,22,82,99]. We found that NF-L (Fig. 4A and 4B), NF-H and phosphorylated NF-H/M (Supplementary Fig. 4A and 4B) are clustered in neurons in the hippocampus of 8.5-month-old *Tmem106b^-/-^* PS19 mice compared to the mice of other genotypes, and co-localized with hTau (Fig. 4A and 4B) (Supplementary Fig. 4A and 4B). Interestingly, we detected a slight but significant decrease in the level of NF-L (but not other NFs) in hippocampal lysates from 6-month-old *Tmem106b^-/-^* mice compared to the WT control (Fig. 4C and 4D). In addition, we also observed a significant accumulation of NF-L in motor neurons in the spinal cord of 8.5-month-old *Tmem106b^-/-^* PS19 mice compared to PS19 mice (Supplementary Fig. 4C and 4D). While the causes of these changes remain to be determined, it suggests that TMEM106B might influence NF dynamics.

**Figure 4.**
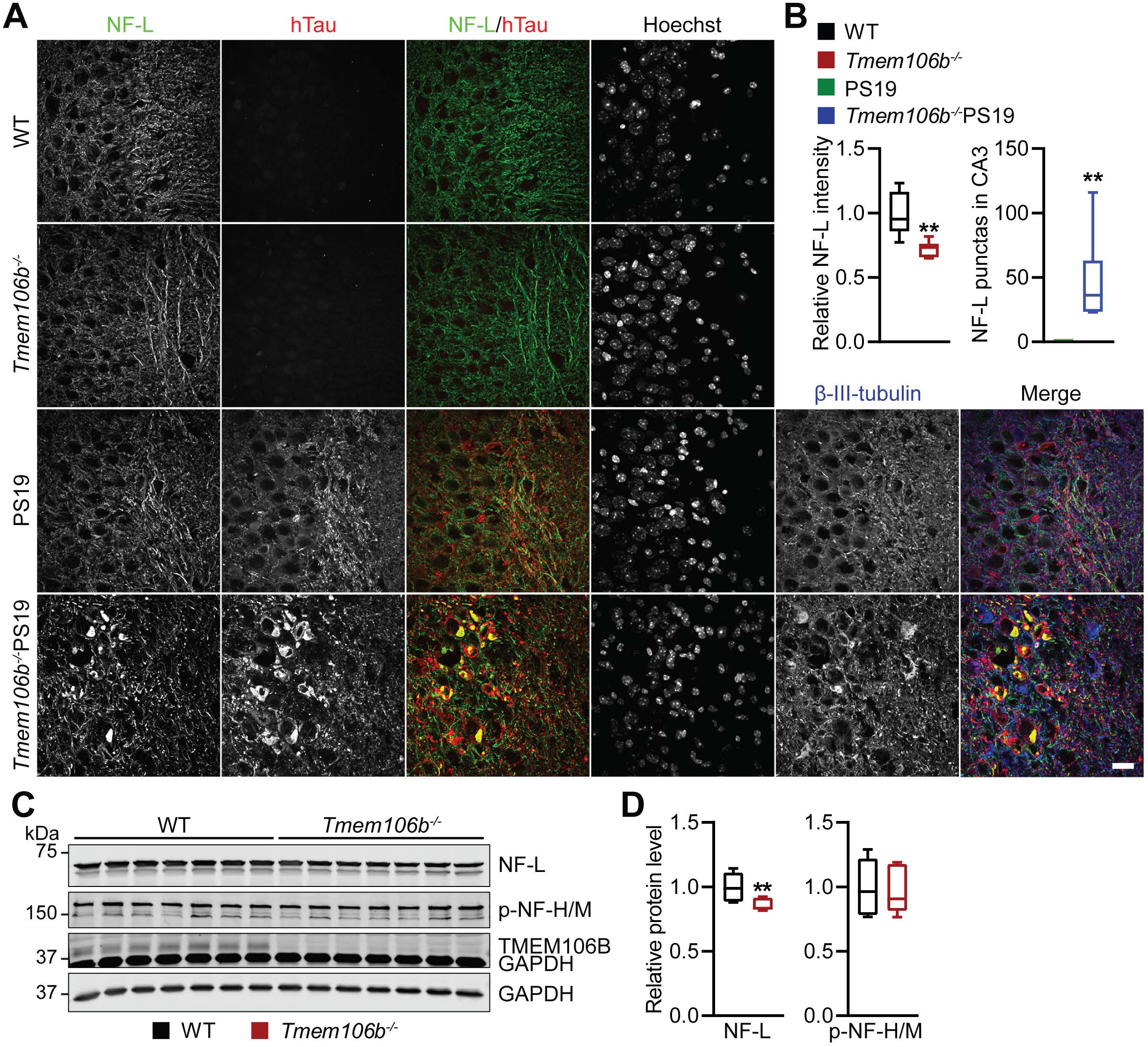
TMEM106B depletion leads to the accumulation of abnormal NF-containing aggregates in PS19 mice. **(A, B)** Representative images of NF-L, hTau, and β-III-tubulin co-immunostainings in CA3 region in the hippocampus of 8.5-month-old WT, *Tmem106b^-/-^*, PS19, and *Tmem106b^-/-^* PS19 mice brain sections. The relative fluorescent intensity and number of fluorescent puncta of NF-L were quantified in B. n=6. Data presented as mean ± SEM. Unpaired Student’s t-test: **, p<0.01. Scale bar: 20 µm. **(C, D)** Western blot analysis of NF-L, p-NF-H/M, TMEM106B, and GAPDH in RIPA-soluble fraction from brain lysates of 6-month-old WT and *Tmem106b^-/-^* mice. Relative levels of indicated proteins were quantified in D. n=7. Data presented as mean ± SEM. Unpaired Student’s t-test: **, p<0.01.

**Figure 5.**
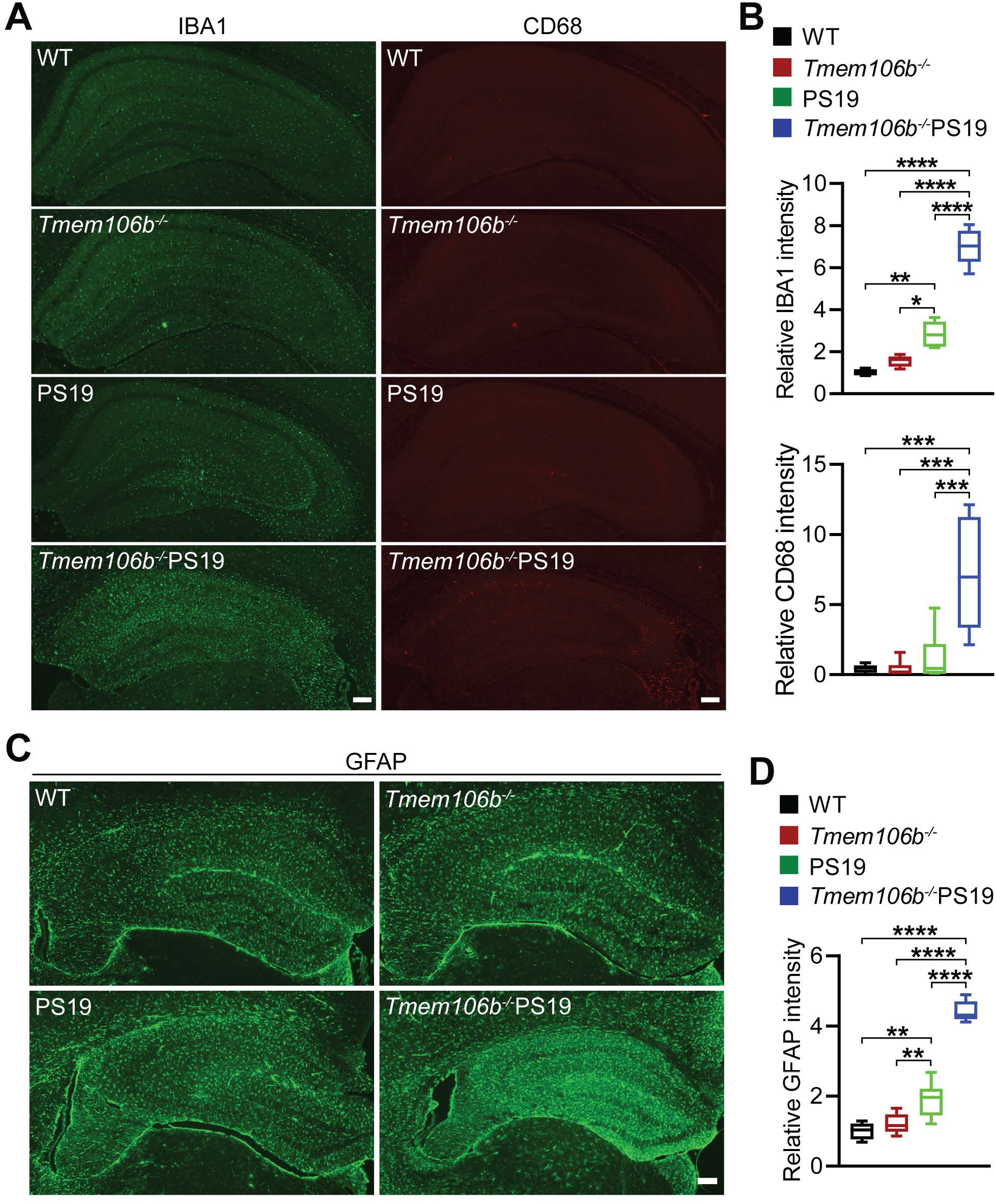
TMEM106B deficiency exacerbates gliosis and neuroinflammation in PS19 mice. (**A, B**) Representative IBA1 and CD68 co-immunostainings in the hippocampus of 8.5-month-old WT, *Tmem106b^-/-^*, PS19, and *Tmem106b^-/-^* PS19 mice. Relative fluorescence intensity of IBA1 or CD68 was quantified in B. n=4-6. Data presented as mean ± SEM. One-way ANOVA tests with Bonferroni’s multiple comparisons: *, p<0.05; **, p<0.01; ***, p<0.001; ****, p<0.0001. Scale bar: 100 µm. (**C, D**) Representative GFAP immunostainings in the hippocampus of 8.5-month-old WT, *Tmem106b^-/-^*, PS19, and *Tmem106b^-/-^* PS19 mice. The relative fluorescence intensity of GFAP was quantified in D. n=5-7. Data presented as mean ± SEM. One-way ANOVA tests with Bonferroni’s multiple comparisons: **, p<0.01; ****, p<0.0001. Scale bar: 100 µm.

### Ablation of TMEM106B enhances gliosis and neuroinflammation in PS19 mice

Tau pathology in the PS19 mice is accompanied by gliosis [96]. To determine whether loss of TMEM106B affects the glia activation in PS19 mice, we performed the immunostaining with antibodies against IBA1, CD68, and GFAP, markers for microglia, active microglia, and astrocyte, respectively. We observed exacerbated microglia and astrocyte activation as shown by increased intensities of IBA1, CD68, and GFAP in the hippocampal sections from 8.5-month-old *Tmem106b^-/-^* PS19 mice compared to the mice with other genotypes (Fig. 5A-5D). Consistently, western blot analysis also showed a significant increase in the protein levels of GFAP in sarkosyl-soluble fraction extracted from the brain of 8.5-month-old *Tmem106b^-/-^* PS19 mice (Supplementary Fig. 5A and 5B).

Since increased microglia and astrocyte activation were detected at 3 and 6 months old of PS19 mice, respectively [96], we also examined the microglia and astrocyte activation in 5 to 5.4-month-old *Tmem106b^-/-^* PS19 mice. No obvious increase in microglia activation was detected in the 5 to 5.4-month-old *Tmem106b^-/-^* PS19 mice compared to PS19 mice (Supplementary Fig. 6), indicating that elevated glia activation seen in 8.5 month-old *Tmem106b^-/-^* PS19 mice might be due to other alterations.

### TMEM106B ablation accelerates autophagy-lysosomal defects in neurons and lysosomal trafficking defects along axons in PS19 mice

TMEM106B is a lysosomal transmembrane protein important for several aspects of lysosomal functions [26]. TMEM106B deficiency in mice leads to autophagic-lysosomal dysfunction [29,53]. To investigate alterations in autophagy lysosomal pathways in *Tmem106b^-/-^* PS19 mice, we first examined the accumulation of lipofuscin, an indicator of lysosomal dysfunction. We found a dramatic increase of lipofuscin signals in the hippocampal CA3 region of *Tmem106b^-/-^* PS19 mice, compared to a subtle accumulation of lipofuscin signal in the PS19 mice and no changes in lipofuscin signal in *Tmem106b^-/-^* mice (Fig. 6A and 6B), indicating that TMEM106B deficiency combined with Tauopathy causes severe lysosomal defects. Next, we examined the protein levels of lysosomal enzymes by western blot analysis. Consistent with the previous studies, the protein levels of lysosomal enzymes Cathepsin D (CathD) and Cathepsin L (CathL) were slightly increased in the brain lysates of *Tmem106b^-/-^*mice compared to the WT mice [29], but it was not altered in PS19 mice (Supplementary Fig. 7A and 7B). Levels of CathD and CathL are not further increased in the brain lysates of *Tmem106b^-/-^* PS19 mice compared to *Tmem106b^-/-^* mice (Supplementary Fig. 7A and 7B). To further determine any changes in lysosomal morphology in neurons, we co-stained brain sections with CathD and NeuN antibodies. Interestingly, we detected a significant reduction of CathD intensities in the hippocampal CA3 pyramidal neurons in 8.5-month-old *Tmem106b^-/-^* PS19 mice compared to PS19 mice (Fig. 6C and 6D). TMEM106B deficiency is known to affect lysosomal trafficking along axons, causing the accumulation of lysosomal vacuoles at the distal end of the axon initial segment (AIS) in motor neurons and Purkinje cells [27,28,53]. We found that CathD-positive vesicles start to accumulate in the axon initial segment (AIS) of Purkinje cells in the cerebellum of 5 to 5.4-month-old *Tmem106b^-/-^* PS19 mice, while PS19, *Tmem106b^-/-^* and WT mice do not show any obvious abnormalities at this age(Fig. 6E and 6F), indicating that loss of TMEM106B in PS19 mice likely results in exacerbated lysosomal trafficking defects and/or other abnormalities in the AIS. Moreover, we observed an accumulation of large CathD-positive clusters out of NeuN-positive neurons in the hippocampus of *Tmem106b^-/-^* PS19 mice, indicating upregulation of CathD in glial cells (Fig. 6C).

**Figure 6.**
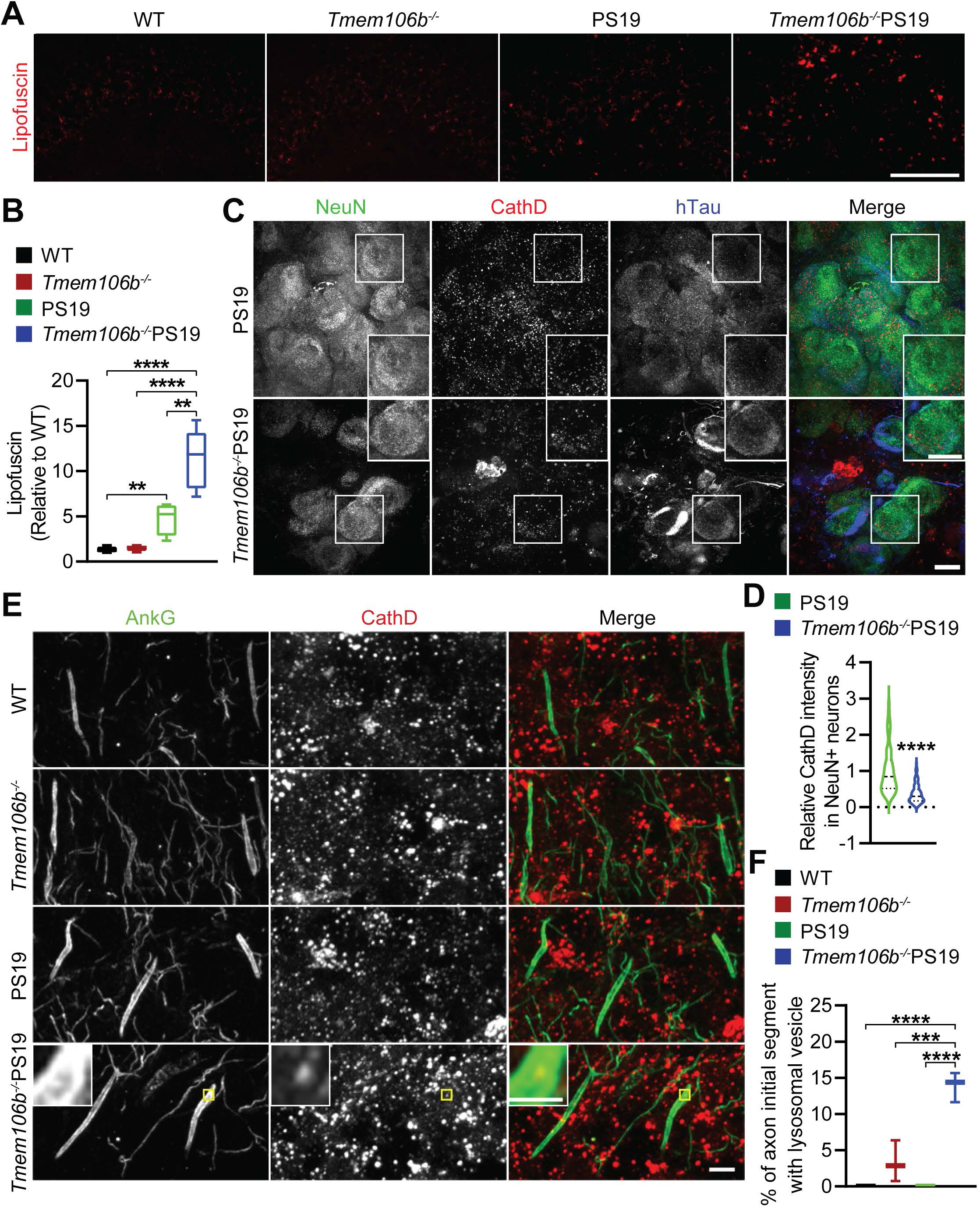
TMEM106B deficiency results in increased lipofuscin accumulation and lysosomal alterations in PS19 mice. **(A, B)** Representative image of lipofuscin autofluorescence in the hippocampal CA3 region of 8.5-month-old WT, *Tmem106b^-/-^*, PS19, and *Tmem106b^-/-^* PS19 mice. The autofluorescence intensity was quantified in B. n=4-6. Data presented as mean ± SEM. One-way ANOVA tests with Bonferroni’s multiple comparisons: **, p<0.01; ****, p<0.0001. Scale bar = 100 µm. **(C, D)** Representative image of NeuN, CathD and hTau co-immunostaining in the hippocampus of 8.5-month-old WT, *Tmem106b^-/-^*, PS19, and *Tmem106b^-/-^* PS19 mice. Relative CathD intensity in NeuN-positive neurons was quantified in D. n=3. Number of neurons =121-125. Data presented as mean ± SEM. Unpaired Student’s t-test: ***, p<0.001. Scale bar: 10 µm. **(E, F)** Representative image of CathD and AnkG co-immunostaining in cerebellum sections from 5 to 5.4-month-old WT, *Tmem106b^-/-^*, PS19 and *Tmem106b^-/-^* PS19 mice. The percentage of axon initial segments with CathD-positive vesicles was quantified in F. n=3. Data presented as mean ± SEM. Unpaired Student’s t-test: *, p<0.05. Scale bar: 1 µm. For Zoom-in images, Scale bar: 0.5 µm.

To further explore alterations in autophagy-lysosome pathway in *Tmem106b^-/-^* PS19 mice, we examined the protein levels of p62, a ubiquitinated cargo receptor for selective autophagy [44,68], and ubiquitinated proteins. Consistent with the previous study [67], the protein level of p62 is reduced in the sarkosyl-soluble fraction from the 8.5-month-old PS19 mouse brain compared to WT mice. However, this decrease was not observed in *Tmem106b^-/-^* PS19 mice (Fig. 7A and 7B). p62 level was not altered in the sarkosyl-insoluble fraction in all the mice analyzed (Fig. 7A and 7B). The protein levels of ubiquitinated proteins are dramatically increased in the sarkosyl-insoluble fraction from 8.5-month-old *Tmem106b^-/-^* PS19 mouse brain when compared to PS19 mice (Fig. 7A and 7B), with slight upregulation seen in PS19 mice compared to WT mice, but no changes are detected in sarkosyl-soluble fractions(Fig. 7A and 7B). Moreover, we observed that both p62 aggregates and ubiquitin-positive inclusions co-localize with hTau in the soma of the pyramidal cell layer in the hippocampus of 8.5-month-old *Tmem106b^-/-^* PS19 mice (Fig. 7C).

**Figure 7.**
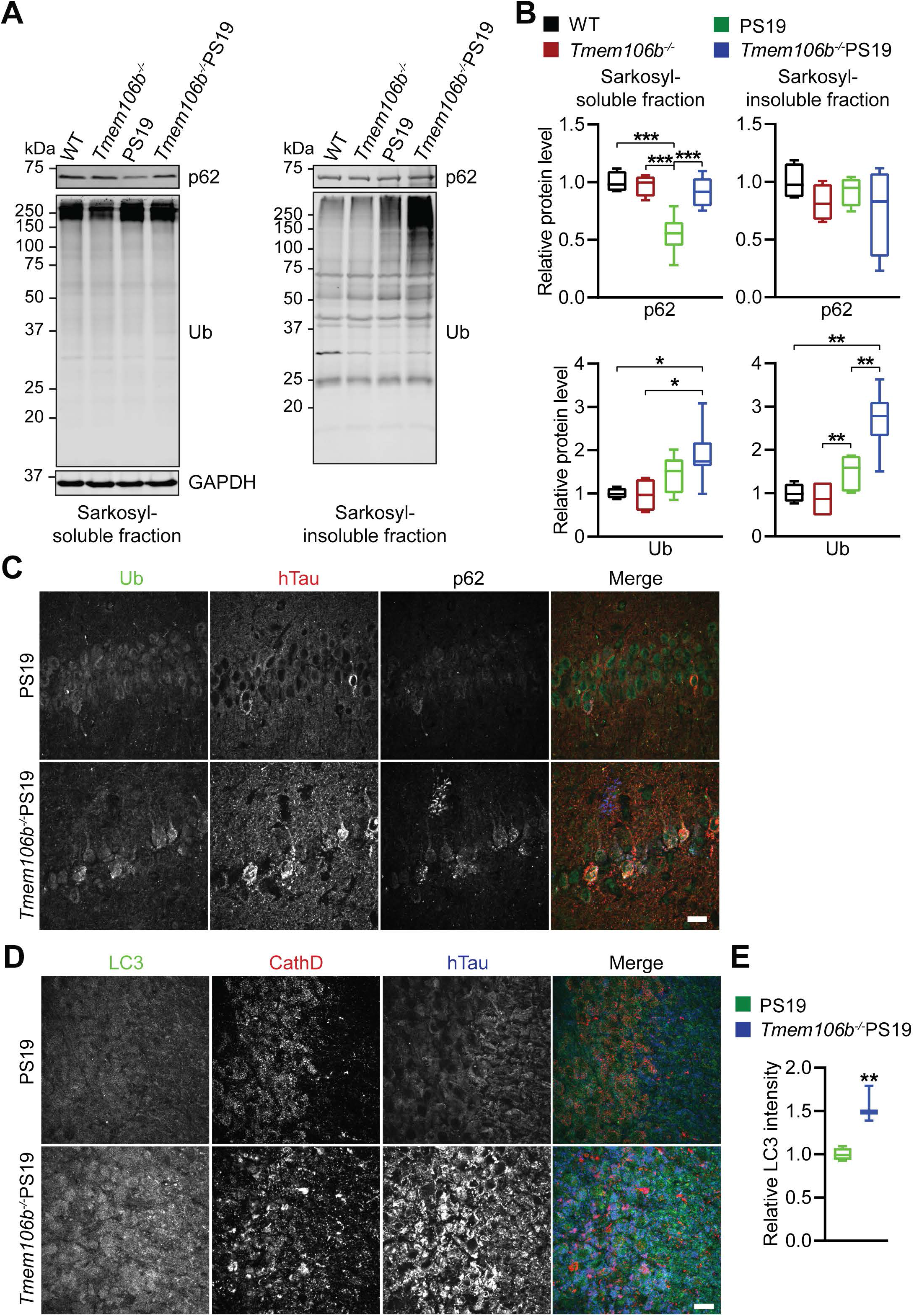
Genetic deletion of TMEM106B upregulated the level of p62 and ubiquitinated proteins in PS19 mice. **(A, B)** Western blot analysis of p62 and Ub in sarkosyl-soluble fraction and sarkosyl-insoluble fraction extracted the brain of 8.5-month-old WT, *Tmem106b^-/-^*, PS19, and *Tmem106b^-/-^* PS19 mice. GAPDH was used as the internal control in the sarkosyl-soluble fraction. The relative level of indicated proteins was quantified in B. n=4-6. Data presented as mean ± SEM. One-way ANOVA tests with Bonferroni’s multiple comparisons: *, p<0.05; **, p<0.01; ***, p<0.001. **(C)** Representative image of Ub, p62, and hTau co-immunostaining in CA1 regions in the hippocampus of 8.5-month-old PS19 and *Tmem106b^-/-^* PS19 mice. Scale bar: 20 µm. **(D, E)** Representative image of LC3 and hTau co-immunostaining in CA3 regions in the hippocampus of 8.5-month-old PS19 and *Tmem106b^-/-^* PS19 mice. The relative fluorescence intensity of LC3 in CA3 somas was quantified in E. n=3-4. Data presented as mean ± SEM. Unpaired Student’s t-test: **, p<0.01. Scale bar: 20 µm.

Finally, we performed the co-immunofluorescence staining using antibodies against LC3, CathD, and hTau. Interestingly, we observed a significant increase of LC3 signals accompanied by the accumulation of hTau in CA3 neuronal soma in 8.5-month-old *Tmem106b^-/-^* PS19 mice compared to PS19 mice, but the the increased LC3 signals were not colocalized with CathD in CA3 neuronal soma (Fig. 7D and 7E),

### TMEM106B ablation upregulates lysosomal CathD in microglia through TFE3/TFEB in PS19 mice

To further determine the sources of the CathD-positive clusters in the CA3 region (Fig. 6C), we co-stained brain sections with antibodies against CathD and IBAI, a microglia marker, or CathD and GFAP, an astrocyte marker. We found that the accumulated CathD-positive clusters co-localize with microglia marker IBA1 (Fig. 8A and 8B), but not astrocyte marker GFAP (Supplementary Fig. 7C), indicating a significant upregulation of CathD in microglia in *Tmem106b^-/-^* PS19 mice.

**Figure 8.**
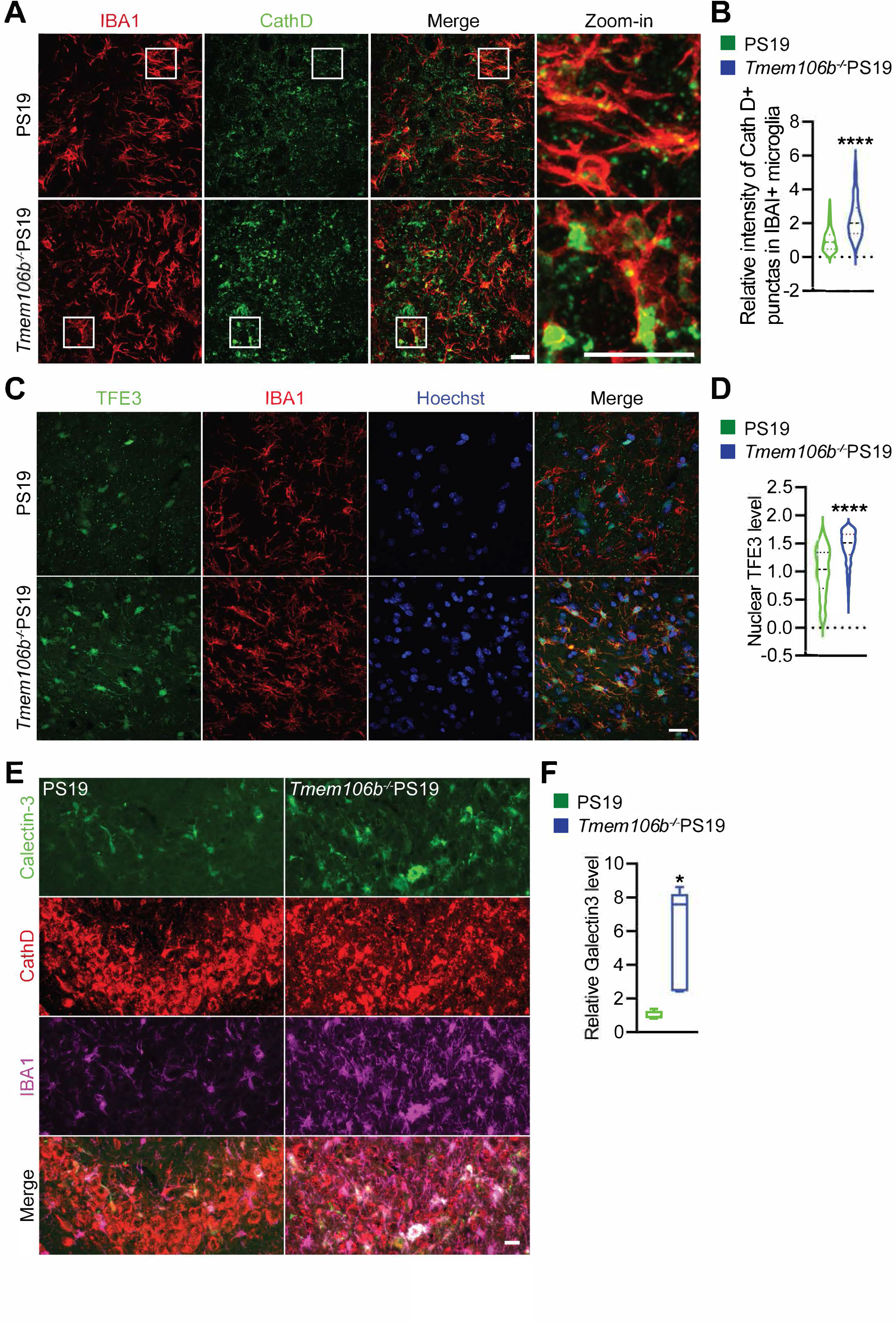
TMEM106B deficiency increases TFE3 nuclear translocation and Galectin 3 upregulation in microglia in PS19 mice. **(A, B)** Representative image of CathD and IBA1 co-immunostaining in CA3 region in the hippocampus of 8.5-month-old PS19 and *Tmem106b^-/-^* PS19 mice. Relative fluorescence intensity of CathD in IBA1-positive microglia was quantified in B. n=3. Data presented as mean ± SEM. Unpaired Student’s t-test: ****, p<0.0001. Scale bar: 20 µm. **(C, D)** Representative TFE3 and IBA1 co-immunostaining in CA3 regions in the hippocampus of 8.5-month-old PS19 and *Tmem106b^-/-^* PS19 mice. Nuclear TFE3 intensity was quantified in D. n=4. Number of cells=311-635. Data presented as mean ± SEM. Unpaired Student’s t-test: ****, p<0.0001. Scale bar: 20 µm. **(E, F)** Representative Galectin-3 and IBA1 co-immunostaining in CA3 regions in the hippocampus of 8.5-month-old PS19 and *Tmem106b^-/-^* PS19 mice. The fluorescence intensity of Galectin-3 in IBA1-positive microglia was quantified in F. n=4-5. Data presented as mean ± SEM. Unpaired Student’s t-test: *, p<0.05. Scale bar: 20 µm.

TFE3, a member of the melanocyte-inducing transcription factor (MiTF) family, is a master regulator in the transcriptional response to starvation or lysosomal stress, and positively upregulates the expression of many genes in the autophagy–lysosome pathway after its translocation into the nucleus [62]. In our previous study, we found that loss of TMEM106B in progranulin-deficient mice results in translocation of TFE3 into the nucleus and the upregulation of lysosomal genes in the microglia [28]. To determine whether TFE3 is activated in the microglia of *Tmem106b^-/-^* PS19 mice, we performed immunostaining using antibodies against TFE3 and IBA1, and observed a drastic increase of nuclear TFE3 signals in the IBA1-positive microglia in the hippocampus of *Tmem106b^-/-^* PS19 mice, compared to that in the PS19 mice (Fig. 8C and 8D). This result suggests that loss of TMEM106B in PS19 mice results in the upregulation of lysosomal genes through TFE3/TFEB transcriptional factors.

Galectin-3 (Gal3), a lectin recognizing β-galactoside, is mainly expressed by activated microglia in the brain [33,40]. Co-immunostaining using antibodies against Galectin-3 and IBA1 shows a significant upregulation of Galectin-3 in the IBA1-positive microglia in *Tmem106b^-/-^* PS19 mice, compared to that in the PS19 mice (Fig. 8E and 8F), indicating alterations in microglial activation in *Tmem106b^-/-^* PS19 mice.

### Loss of TMEM106B does not affect the uptake of mutant hTau in microglia and astrocytes

Microglia, the resident immune cells of the central nervous system (CNS), play crucial roles in neuroinflammation and many other brain disorders [61,74]. Tau pathology induces microglia activation, which further exacerbates Tau pathology and neurodegeneration, indicating a toxic positive feedback loop between Tau pathology and neuroinflammation [7,57,60,78,93]. Since microglia exhibit severe lysosomal defects in microglia in *Tmem106b^-/-^* PS19 mice, we speculated that loss of TMEM106B might affect the uptake of hTau. However, we did not observe a significant difference in Tau signals between PS19 and *Tmem106b^-/-^* PS19 microglia in brain sections (Supplementary Fig. 8A) and the uptake of hTau between WT and *Tmem106b^-/-^* microglia cultured in vitro (Supplementary Fig. 8B). Similarly, we fail to detect any alterations in hTau signals in astrocytes in *Tmem106b^-/-^* PS19 mice (Supplementary Fig. 8C), indicating that the increased Tau pathology in the TMEM106B deficient background might not be due to alterations in astrocyte and microglia activities.

## Discussion

In the present study, we investigate the role of TMEM106B in Tau pathologies using the P301S Tau (PS19) transgenic mice. We demonstrate that loss of TMEM106B exacerbates Tauopathy and its related phenotypes, including (1) accumulation of pathological Tau; (2) neuronal loss and brain atrophy (3) disruption of neuronal cytoskeleton, including microtubules and neurofilaments; (4) gliosis; (5) autophagy-lysosomal defects and lysosomal trafficking along axons. These findings reveal a critical role of TMEM106B in the pathogenesis of Tauopathies and provide new insights into TMEM106B functions in neurodegenerative diseases.

### Loss of TMEM106B exacerbates Tauopathy and neurodegeneration in PS19 mice

TMEM106B risk alleles have been associated with tauopathy in CTE and FTLD-TDP patients [17,52], implying that TMEM106B might influence Tau pathology and Tau-mediated neurodegeneration. In this study we found that loss of TMEM106B dramatically increases Tau accumulation and phosphorylation in PS19 mice (Supplementary Fig. 2A and 2B)(Fig. 2A-2D), resulting in neuronal apoptosis, degeneration (Supplementary Fig. 1D and 1E) and brain atrophy – (Fig. 1). These findings support TMEM106B dysfunction as a strong modifier of Tau pathology and Tau-mediated neurodegeneration in PS19 mice. However, we found that TMEM106B deficiency does not lead to aggregation or phosphorylation of endogenous mouse tau in the PS19 mice (Supplementary Fig. 3), indicating that TMEM106B might specifically affect the behavior of mutant human 4R Tau.

### TMEM106B regulates the cytoskeleton and Tau trafficking

Notably, we found that loss of TMEM106B leads to the redistribution of the pathological hTau and phosphorylated Tau (Ser-404) from axons to soma in PS19 mice at 8.5 months of age (Fig. 2C, 2E and 2F), indicating TMEM106B might affect Tau trafficking. However, this phenotype is not observed in 5 to 5.4-month-old *Tmem106b^-/-^* PS19 mice (Supplementary Fig. 2C), suggesting the redistribution of Tau might be secondary to other earlier alterations in these mice.

Microtubule dysfunction causes protein trafficking defects along axons [6,13,38]. In PS19 mice, a progressive alteration of microtubule accompanied by an early defect in axonal transport in the hippocampus, followed by axonal degeneration has been reported [96]. Consistent with this, we found the accumulation of abnormal β-III-tubulin punctas, which partially colocalized with hTau in hippocampal CA3 region of 8.5-month-old *Tmem106b^-/-^* PS19 mice (Fig. 3A, 4A), indicating a drastic change of microtubule structure in these mice. Acetylation of α-tubulin regulates microtubule dynamics and stability and was impaired in neurons containing Tau neurofibrillary tangles in Alzheimer’s disease [23,39,73]. Loss of microtubule acetylation disrupts axonal transport [18,24,65]. In this study, we found a slight but significant reduction in Ac-tubulin levels in sarkosyl-soluble fraction and a drastic increase of its levels in sarkosyl-insoluble fraction extracted from 5 to 5.4-month-old *Tmem106b^-/-^* PS19 mice (Fig. 3B and 3C). Interestingly, we also observed a slight, but significant decrease of Ac-tubulin in the brain of aged *Tmem106b^-/-^* mice (Fig. 4E and 4F). These findings suggest that loss of TMEM106B alters α-tubulin acetylation during aging and exacerbates acetylation of α-tubulin defects in PS19 mice. Additionally, loss of TMEM106B also exacerbates hyperphosphorylation of hTau, and hyperphosphorylated hTau is known to dissociate from the microtubule, which further interferes with normal microtubule structure and functions, resulting in Tau trafficking defects seen at a later stage.

Disruption of microtubule structure in *Tmem106b^-/-^* PS19 mice could also affect the trafficking of other proteins along axons. It has been reported that neurofilaments are synthesized in the neuronal soma and transported along the axon via slow axonal transport [91]. Interestingly, we observed the accumulation of NF aggregates colocalized with hTau in neuronal soma in the hippocampus and motor neurons in the spinal cord of 8.5-month-old *Tmem106b^-/-^* PS19 mice (Fig. 4, Supplementary Fig. 4), accompanied by the loss of axon in mossy fibers (Fig.1). Additionally, we observed a mild, but significant decrease of NF-L protein in the hippocampus of *Tmem106b^-/-^* mice at 5 months of age (Fig. 4C and 4D). While it remains to be investigated how TMEM106B affects NF-L protein levels and whether TMEM106B affects the axonal trafficking of NF-L, one possibility is that alterations in microtubule structure caused by TMEM106B deficiency lead to axonal trafficking defects of NF, downregulation of NF-L levels and the co-aggregation of NF and hTau in neuronal soma in the PS19 mice.

### Disruption of autophagy lysosome pathway in *Tmem106b^-/-^* PS19 mice

TMEM106B regulates several lysosomal activities, including lysosomal morphology [11,15,46,84], lysosome pH [15,43,45], lysosomal enzyme activities [43,53], lysosome exocytosis [45], lysosomal positioning [29], lysosomal trafficking in dendrites [80] and axons [27,28,53]. It has been demonstrated that phosphorylated Tau and Tau aggregates are degraded by autophagy-lysosome pathway [41,48,59,92] and hyperphosphorylated Tau co-localizes with the autophagosome markers LC3 and p62 proteins in human patients with Tauopathy [71]. In this study, we found that the levels of lysosomal enzyme CathD were reduced in CA3 neurons (Fig. 6C and 6D), but significantly increased in microglia in 8.5-month-old *Tmem106b^-/-^* PS19 mice compared to PS19 mice (Fig. 8A and 8B), indicating alterations in lysosomal activities in these cells. Consistently, we found increased accumulation of p62 and ubiquitinated proteins in the brain of *Tmem106b^-/-^* PS19 mice and co-localization between hTau aggregates, p62 and ubiquitinated proteins (Fig. 7A-C), indicating that loss of TMEM106B results in impaired degradation of hTau and ubiquitinated substrates via autophagy-lysosome pathway.

Hyperphosphorylated and aggregated Tau is known to cause lysosome dysfunction [51,66,100] and defects of lysosomal trafficking along exons [41]. We observed significant accumulation of the large CathD-positive vesicle in the axon initial segment (AIS) of Purkinje cells in the cerebellum of 5.4-month-old *Tmem106b^-/-^* PS19 mice (Fig. 6E and 6F). This phenotype is not observed in age-matched *Tmem106b^-/-^* mice, indicating a synergistic effect of Tau aggregation and TMEM106B deficiency on lysosomal trafficking along axons. [41]. Since loss of TMEM106B in PS19 mice exacerbates Tau pathology (Fig. 2), this could lead to a positive feedback loop resulting in increased lysosomal dysfunction and more Tau aggregation.

### Alterations in microglial activities in *Tmem106b^-/-^* PS19 mice

Studies have shown that microglia internalize and degrade pathological tau in both human AD brain P301S tauopathy mice [9,41,54]. Mutations in triggering receptor expressed on myeloid cell 2 (*TREM2*) gene increase the risk of developing AD dementia and promote tau pathology [4,35,37,49,56,102]. In our recent study, we show that loss of TMEM106B in microglia results in increased lysosomal pH and decreased levels of TREM2, causing decreased microglia survival in response to demyelination [101]. Interestingly, although we observed a significant upregulation of lysosomal CathD possibly due to increased nuclear translocation of TFE3/TFEB in activated microglia in *Tmem106b^-/-^* PS19 mice (Fig. 8A-8D), loss of TMEM106B does not affect the mutant hTau signals in microglia in vivo (Supplementary Fig. 8A) and the uptake of mutant hTau in microglia in vitro (Supplementary Fig. 8B). Moreover, increased Tau aggregation is observed in the hippocampus of 5 to 5.4-month-old *Tmem106b^-/-^* PS19 mice compared to PS19 mice (Supplementary Fig. 2A and 2B), whereas microglia does not appear to show any obvious differences between PS19 and *Tmem106b^-/-^* PS19 mice at this stage (Supplementary Fig. 6), suggesting that increased microglia activation in 8.5 month-old *Tmem106b^-/-^* PS19 mice might be a secondary response to tau aggregation and other earlier events.

Taken together, our study demonstrates TMEM106B loss of function strongly modifies Tauopathy and its associated phenotypes, such as neuroinflammation, disruption of neuronal cytoskeleton, autophagy-lysosomal defects, and subsequent neurodegenerative phenotypes, and provides new insights into the pathophysiological functions of TMEM106B in the brain.

## Material and Methods

### Primary Antibodies and Reagents

The following antibodies were used in this study: mouse anti-GAPDH (Proteintech Group, 60004-1-Ig), rat anti-human Tau (BioLegend, A16103A), Phospho-Tau (Ser404) (Cell signaling, 20194S), Phospho-Tau (Thr205) (Cell signaling, 49561S), mouse anti-Tau (Proteintech Group, 66499-1-Ig), mouse anti-NeuN (Millipore, MAB377), rabbit anti-NeuN (GeneTex, GTX132974), rabbit anti-Cleaved Caspase-3 (Asp175) (Cell signaling, 9661), mouse anti-β-III-tubulin (Promega, G7121), mouse anti-acetylated alpha Tubulin Antibody (6-11B-1) (Santa Cruz, sc-23950), mouse anti-acetylated Tubulin (Lys40) (Proteintech Group, 66200-1-Ig), rabbit anti-alpha-tubulin (Proteintech Group, 11224-1-AP), rabbit anti-Neurofilament-L (C28E10) (Cell signaling, 2837), mouse anti-phosphorylated Neurofilament H & M (NF-H/NF-M) (Biolegend, 837703), chicken anti-Neurofilament H (Encor Biotechnology, CPCA-NF-H), mouse anti-GFAP (GA5) (Cell signaling, 3670S), rabbit anti-IBA-1 (Wako, 01919741), goat anti-IBA1 (Novus Biologicals, NB100-1028), rat anti-CD68 (Bio-Rad, MCA1957), goat anti-CathD (R&D Systems, AF1029), goat anti-CathL (R&D Systems, AF1515), mouse anti-Ankyrin-G clone N106/36 (NeuroMab, N106/36), mouse anti-Galectin-3 (BioLegend, 126702), rabbit anti-TFE3 (Sigma, HPA023881), Rabbit anti-LC3 (Cell signaling, 2775), rabbit anti-p62 (MBL, PM045), mouse anti-Ubiquitin (BioLegend, 646302), and home-made Rabbit anti-TMEM106B antibodies as characterized previously [11].

The following reagents were also used in the study: DMEM (10-017-CV; Cellgro), HBSS (21-020-CV; Cellgro), DMEM/Ham’s F-12 (DMEM/F-12) (10-092-CV; Cellgro), Fetal bovine serum (FBS) (F0926, Sigma), 0.25% Trypsin (25-053-CI; Corning), Pierce BCA Protein Assay Kit (Thermo scientific, 23225), protease inhibitor (Roche, 05056489001), O.C.T compound (Electron Microscopy Sciences, 62550-01), TrueBlack Lipofuscin Autofluorescence Quencher (Biotium, 23007), Odyssey blocking buffer (LI-COR Biosciences, 927-40000), and Fluoromount-G (Thermo scientific, E113391). DyLight 488-Tau P301L oligomer is a kind gift from the late Professor Chad Dickey (University of South Florida).

### Mouse Strains

TMEM106B knockout mice were generated using the CRISPR/Cas9 gene editing technique as described previously [29]. C57/BL6 and B6;C3-Tg(Prnp-MAPT*PS19)PS19Vle/J (Strain #: 008169) mice were obtained from The Jackson Laboratory. All animals (1-6 adult mice per cage) were housed in a 12h light/dark cycle. Only male mice were used for this study. All the mice were housed in the Weill Hall animal facility at Cornell. All animal experiments and procedures were performed according to NIH guidelines and were approved by the Institutional Animal Care and Use Committee at Cornell.

### Cell culture and biochemical assays

Primary microglia were isolated from postnatal 0 (P0) WT and *Tmem106b^-/-^* mice pups, and cultured in DMEM medium with 10% FBS, and 25 U/mL of penicillin/streptomycin. After 12 days of mixed glial cultures, primary microglia are mechanically isolated from the mixed culture by shaking at 200 rpm for 1 hour at 37°C and cultured in DMEM medium with 10% FBS, and 25 U/mL of penicillin/streptomycin for 2 days before experiments.

For Tau P301L oligomer treatment assays, 100 nM DyLight 488-Tau P301L oligomer was added to the medium of primary cultured microglia for 24 hours before fixation.

### Tau extraction

Mice were euthanized with 5% isoflurane and perfused with 1× PBS. The brains were removed and bisected. Half of the brain was snap-frozen with liquid nitrogen and kept at −80°C. The extraction of sarkosyl-insoluble Tau was based on previously published procedures [36]. Briefly, frozen tissues were homogenized on ice with bead homogenizer (Moni International) in 10 volumes of ice-cold H buffer containing 0.8 M NaCl, 10 mM Tris-HCl (pH 7.5), 1 mM EGTA, 10% sucrose, 1 mM PMSF, proteinase and phosphatase inhibitors. After centrifugation at 20,000 × g for 20 minutes at 4□, supernatants were collected, and the pellet was homogenized in 3 vol of H buffer centrifugation at 20,000 × g for 20 minutes. The supernatants were combined and incubated with 1% N-lauroylsarkosynate (sarkosyl) at 37°C for 1 h and ultracentrifuged at 100,000 × g for 1 hour. The supernatants were collected as the sarkosyl-soluble fraction. The pellets were washed with the same buffer and extracted in 5× v/w of Urea buffer (7 M Urea, 2 M Thiourea, 4% CHAPS, 30 mM Tris-HCl, pH 7.5). After sonication, samples were centrifuged at 100,000 × g at 24°C for 1 hour and the supernatant was collected as the sarkosyl-insoluble fraction. Protein concentrations were determined via BCA assay, and then standardized.

### RIPA-soluble fraction preparation

Mice were perfused with 1× PBS and tissues were removed and snap-frozen with liquid nitrogen and kept at −80°C. On the day of the experiment, frozen tissues were thawed and homogenized on ice with bead homogenizer (Moni International) in ice-cold RIPA buffer (150 mM NaCl, 50 mM Tris-HCl (pH 8.0), 1% Triton X-100, 0.5% sodium deoxycholate, 0.1% SDS) with 1 mM PMSF, proteinase and phosphatase inhibitors. After centrifugation at 14,000 × g for 15 minutes at 4°C, supernatants were collected as the RIPA-soluble fraction. Protein concentrations were determined via BCA assay, and then standardized.

### Western blotting

Equal amounts of protein were mixed with an equivalent volume of 2× SDS-sample buffer containing 60 mM Tris-HCl, pH 6.8, 2% SDS, 25% glycerol, 0.1% bromophenol blue, and 5% β-mercaptoethanol (unless mentioned otherwise), and boiled for 5 min. Samples were loaded on 12% home-made polyacrylamide Tris-glycine gels. The gels were initially resolved at 80 V for 15 min. After bromophenol blue ran into separating gel, the gels were resolved at 120 V for ∼1.5 hours, and then transferred onto polyvinylidene difluoride (PVDF) membrane using a wet transfer system with an ice bag at 300 mA for 1 h. The membrane was briefly washed in 1× PBS and blocked in 5% nonfat milk for 1 h, and then incubated with indicated primary antibodies on a rocker overnight at 4°C. The next day, membranes were washed with 1× TBST three times, 10 min each time, and incubated with secondary fluorescent antibodies for 1 h at room temperature. After washing with 1× TBST three times, 10 min each time, the membranes were scanned using an Odyssey Infrared Imaging System (LI-COR Biosciences), and densitometry was performed using Image Studio (LI-COR Biosciences) and ImageJ software.

### Immunofluorescence staining, image acquisition and analysis

For immunostaining, cells were fixed in 4% paraformaldehyde (PFA) and permeabilized with 0.05% saponin or 0.1% Trion X-100 in Odyssey blocking buffer. After washing three times with 1× PBS, cells were incubated with indicated primary antibodies overnight at 4□, following the incubation with secondary fluorescent antibodies at room temperature for 1 hour. Hoechst was used to stain nuclei.

For the brain section staining, mice were euthanized with 5% isoflurane and perfused with 1× PBS. The brain tissues were removed and post-fixed with 4% paraformaldehyde. After dehydration in 30% sucrose buffer, tissues were embedded in O.C.T compound (Electron Microscopy Sciences). 16-µm-thick brain sections were cut with cryotome (Leica). After rehydration in 1× PBS, the brain sections were blocked and permeabilized with either 0.1% saponin in Odyssey blocking buffer or 0.2% Trion X-100 in 1× PBS with 10% horse serum before incubating with indicated primary antibodies overnight at 4□. The next day, sections were washed and then incubated with secondary fluorescent antibodies at room temperature for 1 hour. After washing, the sections were stained with Hoechst, and then mounted using mounting medium (Vector laboratories). Antigen retrieval was performed by microwaving the sections in sodium citrate buffer (pH 6.0) for 10 min. To block the autofluorescence, all sections were incubated with 1× TrueBlack Lipofuscin Autofluorescence Quencher (Biotium) in 70% ethanol for 30 seconds at room temperature before or after the staining process.

Images were acquired on a CSU-X spinning disc confocal microscope (Intelligent Imaging Innovations) with an HQ2 CCD camera (Photometrics) using 40x, 63x and 100x objectives. Five to ten different random images were captured, and the fluorescence intensity was measured directly with ImageJ after a threshold application. Lower magnification images were captured by 4x, 10x, or 20x objectives on a Leica DMi8 inverted microscope. Three to five images were captured from each sample, and the fluorescence intensity was measured directly with ImageJ after a threshold application. Data from ≥4 brains in each genotype were used for quantitative analysis.

### Statistical analysis

All statistical analyses were performed using GraphPad Prism 8. All data are presented as mean ± SEM. Statistical significance was assessed by unpaired Student’s t-test (for two group comparison), and one-way ANOVA tests with Bonferroni’s multiple comparisons (for multiple comparisons). P values less than or equal to 0.05 were considered statistically significant. *p < 0.05; **p < 0.01; ***p < 0.001; ****p < 0.0001.

## Data availability

The data supporting the findings of this study are included in the supplemental material. Additional data are available from the corresponding author on request. No data are deposited in databases.

## Author contributions

T.F. characterized all the phenotypes, analyzed the data and drafted the manuscript. H.D. and T.F. bred the mice. F.H. supervised the project and edited the manuscript. All authors have read and edited the manuscript.

## Acknowledgments

We would like to thank Xiaochun Wu for technical assistance, Dr. Tony Bretscher’s lab for assistance with the confocal microscope. This work is supported by NINDS/NIA (R01NS088448 & R01NS095954) to F.H.

## Ethical Approval and Consent to Participate

All applicable international, national, and/or institutional guidelines for the care and use of animals were followed. The work under animal protocol 2017-0056 is approved by the Institutional Animal Care and Use Committee at Cornell University.

## References

1. Alonso AD, Cohen LS, Corbo C, Morozova V, ElIdrissi A, Phillips G, Kleiman FE (2018) Hyperphosphorylation of Tau Associates With Changes in Its Function Beyond Microtubule Stability. Front Cell Neurosci 12:338. doi:10.3389/fncel.2018.00338

2. Barbier P, Zejneli O, Martinho M, Lasorsa A, Belle V, Smet-Nocca C, Tsvetkov PO, Devred F, Landrieu I (2019) Role of Tau as a Microtubule-Associated Protein: Structural and Functional Aspects. Front Aging Neurosci 11:204. doi:10.3389/fnagi.2019.00204

3. Bejanin A, Schonhaut DR, La Joie R, Kramer JH, Baker SL, Sosa N, Ayakta N, Cantwell A, Janabi M, Lauriola M, O’Neil JP, Gorno-Tempini ML, Miller ZA, Rosen HJ, Miller BL, Jagust WJ, Rabinovici GD (2017) Tau pathology and neurodegeneration contribute to cognitive impairment in Alzheimer’s disease. Brain 140:3286–3300. doi:10.1093/brain/awx243

4. Bemiller SM, McCray TJ, Allan K, Formica SV, Xu G, Wilson G, Kokiko-Cochran ON, Crish SD, Lasagna-Reeves CA, Ransohoff RM, Landreth GE, Lamb BT (2017) TREM2 deficiency exacerbates tau pathology through dysregulated kinase signaling in a mouse model of tauopathy. Mol Neurodegener 12:74. doi:10.1186/s13024-017-0216-6

5. Benear SL, Ngo CT, Olson IR (2020) Dissecting the Fornix in Basic Memory Processes and Neuropsychiatric Disease: A Review. Brain Connect 10:331–354. doi:10.1089/brain.2020.0749

6. Berth SH, Lloyd TE (2023) Disruption of axonal transport in neurodegeneration. J Clin Invest 133. doi:10.1172/JCI168554

7. Bhaskar K, Konerth M, Kokiko-Cochran ON, Cardona A, Ransohoff RM, Lamb BT (2010) Regulation of tau pathology by the microglial fractalkine receptor. Neuron 68:19–31. doi:10.1016/j.neuron.2010.08.023

8. Bieniek KF, Ross OA, Cormier KA, Walton RL, Soto-Ortolaza A, Johnston AE, DeSaro P, Boylan KB, Graff-Radford NR, Wszolek ZK, Rademakers R, Boeve BF, McKee AC, Dickson DW (2015) Chronic traumatic encephalopathy pathology in a neurodegenerative disorders brain bank. Acta Neuropathol 130:877–889. doi:10.1007/s00401-015-1502-4

9. Bolos M, Llorens-Martin M, Jurado-Arjona J, Hernandez F, Rabano A, Avila J (2016) Direct Evidence of Internalization of Tau by Microglia In Vitro and In Vivo. J Alzheimers Dis 50:77–87. doi:10.3233/JAD-150704

10. Bomont P (2021) The dazzling rise of neurofilaments: Physiological functions and roles as biomarkers. Curr Opin Cell Biol 68:181–191. doi:10.1016/j.ceb.2020.10.011

11. Brady OA, Zheng Y, Murphy K, Huang M, Hu F (2013) The frontotemporal lobar degeneration risk factor, TMEM106B, regulates lysosomal morphology and function. Hum Mol Genet 22:685–695. doi:10.1093/hmg/dds475

12. Brandt R, Trushina NI, Bakota L (2020) Much More Than a Cytoskeletal Protein: Physiological and Pathological Functions of the Non-microtubule Binding Region of Tau. Front Neurol 11:590059. doi:10.3389/fneur.2020.590059

13. Buscaglia G, Northington KR, Moore JK, Bates EA (2020) Reduced TUBA1A Tubulin Causes Defects in Trafficking and Impaired Adult Motor Behavior. eNeuro 7. doi:10.1523/ENEURO.0045-20.2020

14. Chang A, Xiang X, Wang J, Lee C, Arakhamia T, Simjanoska M, Wang C, Carlomagno Y, Zhang G, Dhingra S, Thierry M, Perneel J, Heeman B, Forgrave LM, DeTure M, DeMarco ML, Cook CN, Rademakers R, Dickson DW, Petrucelli L, Stowell MHB, Mackenzie IRA, Fitzpatrick AWP (2022) Homotypic fibrillization of TMEM106B across diverse neurodegenerative diseases. Cell 185:1346–1355 e1315. doi:10.1016/j.cell.2022.02.026

15. Chen-Plotkin AS, Unger TL, Gallagher MD, Bill E, Kwong LK, Volpicelli-Daley L, Busch JI, Akle S, Grossman M, Van Deerlin V, Trojanowski JQ, Lee VM (2012) TMEM106B, the risk gene for frontotemporal dementia, is regulated by the microRNA-132/212 cluster and affects progranulin pathways. J Neurosci 32:11213–11227. doi:10.1523/JNEUROSCI.0521-12.2012

16. Chen Y, Yu Y (2023) Tau and neuroinflammation in Alzheimer’s disease: interplay mechanisms and clinical translation. J Neuroinflammation 20:165. doi:10.1186/s12974-023-02853-3

17. Cherry JD, Mez J, Crary JF, Tripodis Y, Alvarez VE, Mahar I, Huber BR, Alosco ML, Nicks R, Abdolmohammadi B, Kiernan PT, Evers L, Svirsky S, Babcock K, Gardner HM, Meng G, Nowinski CJ, Martin BM, Dwyer B, Kowall NW, Cantu RC, Goldstein LE, Katz DI, Stern RA, Farrer LA, McKee AC, Stein TD (2018) Variation in TMEM106B in chronic traumatic encephalopathy. Acta Neuropathol Commun 6:115. doi:10.1186/s40478-018-0619-9

18. Combs B, Mueller RL, Morfini G, Brady ST, Kanaan NM (2019) Tau and Axonal Transport Misregulation in Tauopathies. Adv Exp Med Biol 1184:81–95. doi:10.1007/978-981-32-9358-8_7

19. Copley KE, Shorter J (2022) Flying under the radar: TMEM106B(120-254) fibrils break out in diverse neurodegenerative disorders. Cell 185:1290–1292. doi:10.1016/j.cell.2022.03.032

20. Cruchaga C, Graff C, Chiang HH, Wang J, Hinrichs AL, Spiegel N, Bertelsen S, Mayo K, Norton JB, Morris JC, Goate A (2011) Association of TMEM106B gene polymorphism with age at onset in granulin mutation carriers and plasma granulin protein levels. Arch Neurol 68:581–586. doi:10.1001/archneurol.2010.350

21. Deming Y, Cruchaga C (2014) TMEM106B: a strong FTLD disease modifier. Acta Neuropathol 127:419–422. doi:10.1007/s00401-014-1249-3

22. Dong DL, Xu ZS, Hart GW, Cleveland DW (1996) Cytoplasmic O-GlcNAc modification of the head domain and the KSP repeat motif of the neurofilament protein neurofilament-H. J Biol Chem 271:20845–20852. doi:10.1074/jbc.271.34.20845

23. Eshun-Wilson L, Zhang R, Portran D, Nachury MV, Toso DB, Lohr T, Vendruscolo M, Bonomi M, Fraser JS, Nogales E (2019) Effects of alpha-tubulin acetylation on microtubule structure and stability. Proc Natl Acad Sci U S A 116:10366–10371. doi:10.1073/pnas.1900441116

24. Even A, Morelli G, Broix L, Scaramuzzino C, Turchetto S, Gladwyn-Ng I, Le Bail R, Shilian M, Freeman S, Magiera MM, Jijumon AS, Krusy N, Malgrange B, Brone B, Dietrich P, Dragatsis I, Janke C, Saudou F, Weil M, Nguyen L (2019) ATAT1-enriched vesicles promote microtubule acetylation via axonal transport. Sci Adv 5:eaax2705. doi:10.1126/sciadv.aax2705

25. Fan Y, Zhao Q, Xia W, Tao Y, Yu W, Chen M, Liu Y, Zhao J, Shen Y, Sun Y, Si C, Zhang S, Zhang Y, Li W, Liu C, Wang J, Li D (2022) Generic amyloid fibrillation of TMEM106B in patient with Parkinson’s disease dementia and normal elders. Cell Res 32:585–588. doi:10.1038/s41422-022-00665-3

26. Feng T, Lacrampe A, Hu F (2021) Physiological and pathological functions of TMEM106B: a gene associated with brain aging and multiple brain disorders. Acta Neuropathol 141:327–339. doi:10.1007/s00401-020-02246-3

27. Feng T, Luan L, Katz, II, Ullah M, Van Deerlin VM, Trojanowski JQ, Lee EB, Hu F (2022) TMEM106B deficiency impairs cerebellar myelination and synaptic integrity with Purkinje cell loss. Acta Neuropathol Commun 10:33. doi:10.1186/s40478-022-01334-7

28. Feng T, Mai S, Roscoe JM, Sheng RR, Ullah M, Zhang J, Katz, II, Yu H, Xiong W, Hu F (2020) Loss of TMEM106B and PGRN leads to severe lysosomal abnormalities and neurodegeneration in mice. EMBO Rep 21:e50219. doi:10.15252/embr.202050219

29. Feng T, Sheng RR, Sole-Domenech S, Ullah M, Zhou X, Mendoza CS, Enriquez LCM, Katz, II, Paushter DH, Sullivan PM, Wu X, Maxfield FR, Hu F (2020) A role of the frontotemporal lobar degeneration risk factor TMEM106B in myelination. Brain 143:2255–2271. doi:10.1093/brain/awaa154

30. Finch N, Carrasquillo MM, Baker M, Rutherford NJ, Coppola G, Dejesus-Hernandez M, Crook R, Hunter T, Ghidoni R, Benussi L, Crook J, Finger E, Hantanpaa KJ, Karydas AM, Sengdy P, Gonzalez J, Seeley WW, Johnson N, Beach TG, Mesulam M, Forloni G, Kertesz A, Knopman DS, Uitti R, White CL, 3rd, Caselli R, Lippa C, Bigio EH, Wszolek ZK, Binetti G, Mackenzie IR, Miller BL, Boeve BF, Younkin SG, Dickson DW, Petersen RC, Graff-Radford NR, Geschwind DH, Rademakers R (2011) TMEM106B regulates progranulin levels and the penetrance of FTLD in GRN mutation carriers. Neurology 76:467–474. doi:10.1212/WNL.0b013e31820a0e3b

31. Gafson AR, Barthelemy NR, Bomont P, Carare RO, Durham HD, Julien JP, Kuhle J, Leppert D, Nixon RA, Weller RO, Zetterberg H, Matthews PM (2020) Neurofilaments: neurobiological foundations for biomarker applications. Brain 143:1975–1998. doi:10.1093/brain/awaa098

32. Gallagher MD, Suh E, Grossman M, Elman L, McCluskey L, Van Swieten JC, Al-Sarraj S, Neumann M, Gelpi E, Ghetti B, Rohrer JD, Halliday G, Van Broeckhoven C, Seilhean D, Shaw PJ, Frosch MP, Alafuzoff I, Antonell A, Bogdanovic N, Brooks W, Cairns NJ, Cooper-Knock J, Cotman C, Cras P, Cruts M, De Deyn PP, DeCarli C, Dobson-Stone C, Engelborghs S, Fox N, Galasko D, Gearing M, Gijselinck I, Grafman J, Hartikainen P, Hatanpaa KJ, Highley JR, Hodges J, Hulette C, Ince PG, Jin LW, Kirby J, Kofler J, Kril J, Kwok JB, Levey A, Lieberman A, Llado A, Martin JJ, Masliah E, McDermott CJ, McKee A, McLean C, Mead S, Miller CA, Miller J, Munoz DG, Murrell J, Paulson H, Piguet O, Rossor M, Sanchez-Valle R, Sano M, Schneider J, Silbert LC, Spina S, van der Zee J, Van Langenhove T, Warren J, Wharton SB, White CL, 3rd, Woltjer RL, Trojanowski JQ, Lee VM, Van Deerlin V, Chen-Plotkin AS (2014) TMEM106B is a genetic modifier of frontotemporal lobar degeneration with C9orf72 hexanucleotide repeat expansions. Acta Neuropathol 127:407–418. doi:10.1007/s00401-013-1239-x

33. Garcia-Revilla J, Boza-Serrano A, Espinosa-Oliva AM, Soto MS, Deierborg T, Ruiz R, de Pablos RM, Burguillos MA, Venero JL (2022) Galectin-3, a rising star in modulating microglia activation under conditions of neurodegeneration. Cell Death Dis 13:628. doi:10.1038/s41419-022-05058-3

34. Goedert M, Eisenberg DS, Crowther RA (2017) Propagation of Tau Aggregates and Neurodegeneration. Annu Rev Neurosci 40:189–210. doi:10.1146/annurev-neuro-072116-031153

35. Gratuze M, Chen Y, Parhizkar S, Jain N, Strickland MR, Serrano JR, Colonna M, Ulrich JD, Holtzman DM (2021) Activated microglia mitigate Abeta-associated tau seeding and spreading. J Exp Med 218. doi:10.1084/jem.20210542

36. Greenberg SG, Davies P (1990) A preparation of Alzheimer paired helical filaments that displays distinct tau proteins by polyacrylamide gel electrophoresis. Proc Natl Acad Sci U S A 87:5827–5831. doi:10.1073/pnas.87.15.5827

37. Guerreiro R, Wojtas A, Bras J, Carrasquillo M, Rogaeva E, Majounie E, Cruchaga C, Sassi C, Kauwe JS, Younkin S, Hazrati L, Collinge J, Pocock J, Lashley T, Williams J, Lambert JC, Amouyel P, Goate A, Rademakers R, Morgan K, Powell J, St George-Hyslop P, Singleton A, Hardy J, Alzheimer Genetic Analysis G (2013) TREM2 variants in Alzheimer’s disease. N Engl J Med 368:117–127. doi:10.1056/NEJMoa1211851

38. Guo W, Stoklund Dittlau K, Van Den Bosch L (2020) Axonal transport defects and neurodegeneration: Molecular mechanisms and therapeutic implications. Semin Cell Dev Biol 99:133–150. doi:10.1016/j.semcdb.2019.07.010

39. Hempen B, Brion JP (1996) Reduction of acetylated alpha-tubulin immunoreactivity in neurofibrillary tangle-bearing neurons in Alzheimer’s disease. J Neuropathol Exp Neurol 55:964–972. doi:10.1097/00005072-199609000-00003

40. Jia J, Claude-Taupin A, Gu Y, Choi SW, Peters R, Bissa B, Mudd MH, Allers L, Pallikkuth S, Lidke KA, Salemi M, Phinney B, Mari M, Reggiori F, Deretic V (2020) Galectin-3 Coordinates a Cellular System for Lysosomal Repair and Removal. Dev Cell 52:69–87 e68. doi:10.1016/j.devcel.2019.10.025

41. Jiang S, Bhaskar K (2020) Degradation and Transmission of Tau by Autophagic-Endolysosomal Networks and Potential Therapeutic Targets for Tauopathy. Front Mol Neurosci 13:586731. doi:10.3389/fnmol.2020.586731

42. Jiang YX, Cao Q, Sawaya MR, Abskharon R, Ge P, DeTure M, Dickson DW, Fu JY, Ogorzalek Loo RR, Loo JA, Eisenberg DS (2022) Amyloid fibrils in FTLD-TDP are composed of TMEM106B and not TDP-43. Nature 605:304–309. doi:10.1038/s41586-022-04670-9

43. Klein ZA, Takahashi H, Ma M, Stagi M, Zhou M, Lam TT, Strittmatter SM (2017) Loss of TMEM106B Ameliorates Lysosomal and Frontotemporal Dementia-Related Phenotypes in Progranulin-Deficient Mice. Neuron 95:281–296 e286. doi:10.1016/j.neuron.2017.06.026

44. Komatsu M, Waguri S, Koike M, Sou YS, Ueno T, Hara T, Mizushima N, Iwata J, Ezaki J, Murata S, Hamazaki J, Nishito Y, Iemura S, Natsume T, Yanagawa T, Uwayama J, Warabi E, Yoshida H, Ishii T, Kobayashi A, Yamamoto M, Yue Z, Uchiyama Y, Kominami E, Tanaka K (2007) Homeostatic levels of p62 control cytoplasmic inclusion body formation in autophagy-deficient mice. Cell 131:1149–1163. doi:10.1016/j.cell.2007.10.035

45. Kundu ST, Grzeskowiak CL, Fradette JJ, Gibson LA, Rodriguez LB, Creighton CJ, Scott KL, Gibbons DL (2018) TMEM106B drives lung cancer metastasis by inducing TFEB-dependent lysosome synthesis and secretion of cathepsins. Nat Commun 9:2731. doi:10.1038/s41467-018-05013-x

46. Lang CM, Fellerer K, Schwenk BM, Kuhn PH, Kremmer E, Edbauer D, Capell A, Haass C (2012) Membrane orientation and subcellular localization of transmembrane protein 106B (TMEM106B), a major risk factor for frontotemporal lobar degeneration. J Biol Chem 287:19355–19365. doi:10.1074/jbc.M112.365098

47. Lattante S, Le Ber I, Galimberti D, Serpente M, Rivaud-Pechoux S, Camuzat A, Clot F, Fenoglio C, French research network on FTD, Ftd ALS, Scarpini E, Brice A, Kabashi E (2014) Defining the association of TMEM106B variants among frontotemporal lobar degeneration patients with GRN mutations and C9orf72 repeat expansions. Neurobiol Aging 35:2658 e2651–2658 e2655. doi:10.1016/j.neurobiolaging.2014.06.023

48. Lee MJ, Lee JH, Rubinsztein DC (2013) Tau degradation: the ubiquitin-proteasome system versus the autophagy-lysosome system. Prog Neurobiol 105:49–59. doi:10.1016/j.pneurobio.2013.03.001

49. Leyns CEG, Gratuze M, Narasimhan S, Jain N, Koscal LJ, Jiang H, Manis M, Colonna M, Lee VMY, Ulrich JD, Holtzman DM (2019) TREM2 function impedes tau seeding in neuritic plaques. Nat Neurosci 22:1217–1222. doi:10.1038/s41593-019-0433-0

50. Li ZH, Lu J, Tay SS, Wu YJ, Strong MJ, He BP (2006) Mice with targeted disruption of neurofilament light subunit display formation of protein aggregation in motoneurons and downregulation of complement receptor type 3 alpha subunit in microglia in the spinal cord at their earlier age: a possible feature in pre-clinical development of neurodegenerative diseases. Brain Res 1113:200–209. doi:10.1016/j.brainres.2006.07.041

51. Lim F, Hernandez F, Lucas JJ, Gomez-Ramos P, Moran MA, Avila J (2001) FTDP-17 mutations in tau transgenic mice provoke lysosomal abnormalities and Tau filaments in forebrain. Mol Cell Neurosci 18:702–714. doi:10.1006/mcne.2001.1051

52. Llibre-Guerra JJ, Lee SE, Suemoto CK, Ehrenberg AJ, Kovacs GG, Karydas A, Staffaroni A, Franca Resende EP, Kim EJ, Hwang JH, Ramos EM, Wojta KJ, Pasquini L, Pang SY, Spina S, Allen IE, Kramer J, Miller BL, Seeley WW, Grinberg LT (2021) A novel temporal-predominant neuro-astroglial tauopathy associated with TMEM106B gene polymorphism in FTLD/ALS-TDP. Brain Pathol 31:267–282. doi:10.1111/bpa.12924

53. Luningschror P, Werner G, Stroobants S, Kakuta S, Dombert B, Sinske D, Wanner R, Lullmann-Rauch R, Wefers B, Wurst W, D’Hooge R, Uchiyama Y, Sendtner M, Haass C, Saftig P, Knoll B, Capell A, Damme M (2020) The FTLD Risk Factor TMEM106B Regulates the Transport of Lysosomes at the Axon Initial Segment of Motoneurons. Cell Rep 30:3506–3519 e3506. doi:10.1016/j.celrep.2020.02.060

54. Luo W, Liu W, Hu X, Hanna M, Caravaca A, Paul SM (2015) Microglial internalization and degradation of pathological tau is enhanced by an anti-tau monoclonal antibody. Sci Rep 5:11161. doi:10.1038/srep11161

55. Mandelkow EM, Mandelkow E (2012) Biochemistry and cell biology of tau protein in neurofibrillary degeneration. Cold Spring Harb Perspect Med 2:a006247. doi:10.1101/cshperspect.a006247

56. Mao F, Robinson JL, Unger T, Posavi M, Amado DA, Elman L, Grossman M, Wolk DA, Lee EB, Van Deerlin VM, Porta S, Lee VMY, Trojanowski JQ, Chen-Plotkin AS (2021) TMEM106B modifies TDP-43 pathology in human ALS brain and cell-based models of TDP-43 proteinopathy. Acta Neuropathol 142:629–642. doi:10.1007/s00401-021-02330-2

57. Maphis N, Xu G, Kokiko-Cochran ON, Jiang S, Cardona A, Ransohoff RM, Lamb BT, Bhaskar K (2015) Reactive microglia drive tau pathology and contribute to the spreading of pathological tau in the brain. Brain 138:1738–1755. doi:10.1093/brain/awv081

58. McLean JR, Sanelli TR, Leystra-Lantz C, He BP, Strong MJ (2005) Temporal profiles of neuronal degeneration, glial proliferation, and cell death in hNFL(+/+) and NFL(-/-) mice. Glia 52:59–69. doi:10.1002/glia.20218

59. Menzies FM, Fleming A, Caricasole A, Bento CF, Andrews SP, Ashkenazi A, Fullgrabe J, Jackson A, Jimenez Sanchez M, Karabiyik C, Licitra F, Lopez Ramirez A, Pavel M, Puri C, Renna M, Ricketts T, Schlotawa L, Vicinanza M, Won H, Zhu Y, Skidmore J, Rubinsztein DC (2017) Autophagy and Neurodegeneration: Pathogenic Mechanisms and Therapeutic Opportunities. Neuron 93:1015–1034. doi:10.1016/j.neuron.2017.01.022

60. Morales I, Jimenez JM, Mancilla M, Maccioni RB (2013) Tau oligomers and fibrils induce activation of microglial cells. J Alzheimers Dis 37:849–856. doi:10.3233/JAD-131843

61. Muzio L, Viotti A, Martino G (2021) Microglia in Neuroinflammation and Neurodegeneration: From Understanding to Therapy. Front Neurosci 15:742065. doi:10.3389/fnins.2021.742065

62. Napolitano G, Ballabio A (2016) TFEB at a glance. J Cell Sci 129:2475–2481. doi:10.1242/jcs.146365

63. Nelson PT, Dickson DW, Trojanowski JQ, Jack CR, Boyle PA, Arfanakis K, Rademakers R, Alafuzoff I, Attems J, Brayne C, Coyle-Gilchrist ITS, Chui HC, Fardo DW, Flanagan ME, Halliday G, Hokkanen SRK, Hunter S, Jicha GA, Katsumata Y, Kawas CH, Keene CD, Kovacs GG, Kukull WA, Levey AI, Makkinejad N, Montine TJ, Murayama S, Murray ME, Nag S, Rissman RA, Seeley WW, Sperling RA, White Iii CL, Yu L, Schneider JA (2019) Limbic-predominant age-related TDP-43 encephalopathy (LATE): consensus working group report. Brain 142:1503–1527. doi:10.1093/brain/awz099

64. Nelson PT, Wang WX, Partch AB, Monsell SE, Valladares O, Ellingson SR, Wilfred BR, Naj AC, Wang LS, Kukull WA, Fardo DW (2015) Reassessment of risk genotypes (GRN, TMEM106B, and ABCC9 variants) associated with hippocampal sclerosis of aging pathology. J Neuropathol Exp Neurol 74:75–84. doi:10.1097/NEN.0000000000000151

65. Neumann B, Hilliard MA (2014) Loss of MEC-17 leads to microtubule instability and axonal degeneration. Cell Rep 6:93–103. doi:10.1016/j.celrep.2013.12.004

66. Nixon RA, Cataldo AM, Mathews PM (2000) The endosomal-lysosomal system of neurons in Alzheimer’s disease pathogenesis: a review. Neurochem Res 25:1161–1172. doi:10.1023/a:1007675508413

67. Ono M, Komatsu M, Ji B, Takado Y, Shimojo M, Minamihisamatsu T, Warabi E, Yanagawa T, Matsumoto G, Aoki I, Kanaan NM, Suhara T, Sahara N, Higuchi M (2022) Central role for p62/SQSTM1 in the elimination of toxic tau species in a mouse model of tauopathy. Aging Cell 21:e13615. doi:10.1111/acel.13615

68. Pankiv S, Clausen TH, Lamark T, Brech A, Bruun JA, Outzen H, Overvatn A, Bjorkoy G, Johansen T (2007) p62/SQSTM1 binds directly to Atg8/LC3 to facilitate degradation of ubiquitinated protein aggregates by autophagy. J Biol Chem 282:24131–24145. doi:10.1074/jbc.M702824200

69. Perneel J, Neumann M, Heeman B, Cheung S, Van den Broeck M, Wynants S, Baker M, Vicente CT, Faura J, Rademakers R, Mackenzie IRA (2023) Accumulation of TMEM106B C-terminal fragments in neurodegenerative disease and aging. Acta Neuropathol 145:285–302. doi:10.1007/s00401-022-02531-3

70. Perneel J, Rademakers R (2022) Identification of TMEM106B amyloid fibrils provides an updated view of TMEM106B biology in health and disease. Acta Neuropathol 144:807–819. doi:10.1007/s00401-022-02486-5

71. Piras A, Collin L, Gruninger F, Graff C, Ronnback A (2016) Autophagic and lysosomal defects in human tauopathies: analysis of post-mortem brain from patients with familial Alzheimer disease, corticobasal degeneration and progressive supranuclear palsy. Acta Neuropathol Commun 4:22. doi:10.1186/s40478-016-0292-9

72. Portier MM (1992) [Neuronal cytoskeleton: structural, functional and dynamic aspects]. Rev Neurol (Paris) 148:1–19

73. Portran D, Schaedel L, Xu Z, Thery M, Nachury MV (2017) Tubulin acetylation protects long-lived microtubules against mechanical aging. Nat Cell Biol 19:391–398. doi:10.1038/ncb3481

74. Prinz M, Masuda T, Wheeler MA, Quintana FJ (2021) Microglia and Central Nervous System-Associated Macrophages-From Origin to Disease Modulation. Annu Rev Immunol 39:251–277. doi:10.1146/annurev-immunol-093019-110159

75. Rawat P, Sehar U, Bisht J, Selman A, Culberson J, Reddy PH (2022) Phosphorylated Tau in Alzheimer’s Disease and Other Tauopathies. Int J Mol Sci 23. doi:10.3390/ijms232112841

76. Rhinn H, Abeliovich A (2017) Differential Aging Analysis in Human Cerebral Cortex Identifies Variants in TMEM106B and GRN that Regulate Aging Phenotypes. Cell Syst 4:404–415 e405. doi:10.1016/j.cels.2017.02.009

77. Rutherford NJ, Carrasquillo MM, Li M, Bisceglio G, Menke J, Josephs KA, Parisi JE, Petersen RC, Graff-Radford NR, Younkin SG, Dickson DW, Rademakers R (2012) TMEM106B risk variant is implicated in the pathologic presentation of Alzheimer disease. Neurology 79:717–718. doi:10.1212/WNL.0b013e318264e3ac

78. Samudra N, Lane-Donovan C, VandeVrede L, Boxer AL (2023) Tau pathology in neurodegenerative disease: disease mechanisms and therapeutic avenues. J Clin Invest 133. doi:10.1172/JCI168553

79. Schweighauser M, Arseni D, Bacioglu M, Huang M, Lovestam S, Shi Y, Yang Y, Zhang W, Kotecha A, Garringer HJ, Vidal R, Hallinan GI, Newell KL, Tarutani A, Murayama S, Miyazaki M, Saito Y, Yoshida M, Hasegawa K, Lashley T, Revesz T, Kovacs GG, van Swieten J, Takao M, Hasegawa M, Ghetti B, Spillantini MG, Ryskeldi-Falcon B, Murzin AG, Goedert M, Scheres SHW (2022) Age-dependent formation of TMEM106B amyloid filaments in human brains. Nature 605:310–314. doi:10.1038/s41586-022-04650-z

80. Schwenk BM, Lang CM, Hogl S, Tahirovic S, Orozco D, Rentzsch K, Lichtenthaler SF, Hoogenraad CC, Capell A, Haass C, Edbauer D (2014) The FTLD risk factor TMEM106B and MAP6 control dendritic trafficking of lysosomes. EMBO J 33:450–467. doi:10.1002/embj.201385857

81. Sferra A, Nicita F, Bertini E (2020) Microtubule Dysfunction: A Common Feature of Neurodegenerative Diseases. Int J Mol Sci 21. doi:10.3390/ijms21197354

82. Sihag RK, Inagaki M, Yamaguchi T, Shea TB, Pant HC (2007) Role of phosphorylation on the structural dynamics and function of types III and IV intermediate filaments. Exp Cell Res 313:2098–2109. doi:10.1016/j.yexcr.2007.04.010

83. Simons C, Dyment D, Bent SJ, Crawford J, D’Hooghe M, Kohlschutter A, Venkateswaran S, Helman G, Poll-The BT, Makowski CC, Ito Y, Kernohan K, Hartley T, Waisfisz Q, Taft RJ, van der Knaap MS, Wolf NI (2017) A recurrent de novo mutation in TMEM106B causes hypomyelinating leukodystrophy. Brain 140:3105–3111. doi:10.1093/brain/awx314

84. Stagi M, Klein ZA, Gould TJ, Bewersdorf J, Strittmatter SM (2014) Lysosome size, motility and stress response regulated by fronto-temporal dementia modifier TMEM106B. Mol Cell Neurosci 61:226–240. doi:10.1016/j.mcn.2014.07.006

85. Sun Y, Guo Y, Feng X, Jia M, Ai N, Dong Y, Zheng Y, Fu L, Yu B, Zhang H, Wu J, Yu X, Wu H, Kong W (2020) The behavioural and neuropathologic sexual dimorphism and absence of MIP-3alpha in tau P301S mouse model of Alzheimer’s disease. J Neuroinflammation 17:72. doi:10.1186/s12974-020-01749-w

86. Tropea TF, Mak J, Guo MH, Xie SX, Suh E, Rick J, Siderowf A, Weintraub D, Grossman M, Irwin D, Wolk DA, Trojanowski JQ, Van Deerlin V, Chen-Plotkin AS (2019) TMEM106B Effect on cognition in Parkinson disease and frontotemporal dementia. Ann Neurol 85:801–811. doi:10.1002/ana.25486

87. van Blitterswijk M, Mullen B, Nicholson AM, Bieniek KF, Heckman MG, Baker MC, DeJesus-Hernandez M, Finch NA, Brown PH, Murray ME, Hsiung GY, Stewart H, Karydas AM, Finger E, Kertesz A, Bigio EH, Weintraub S, Mesulam M, Hatanpaa KJ, White CL, 3rd, Strong MJ, Beach TG, Wszolek ZK, Lippa C, Caselli R, Petrucelli L, Josephs KA, Parisi JE, Knopman DS, Petersen RC, Mackenzie IR, Seeley WW, Grinberg LT, Miller BL, Boylan KB, Graff-Radford NR, Boeve BF, Dickson DW, Rademakers R (2014) TMEM106B protects C9ORF72 expansion carriers against frontotemporal dementia. Acta Neuropathol 127:397–406. doi:10.1007/s00401-013-1240-4

88. Van Deerlin VM, Sleiman PM, Martinez-Lage M, Chen-Plotkin A, Wang LS, Graff-Radford NR, Dickson DW, Rademakers R, Boeve BF, Grossman M, Arnold SE, Mann DM, Pickering-Brown SM, Seelaar H, Heutink P, van Swieten JC, Murrell JR, Ghetti B, Spina S, Grafman J, Hodges J, Spillantini MG, Gilman S, Lieberman AP, Kaye JA, Woltjer RL, Bigio EH, Mesulam M, Al-Sarraj S, Troakes C, Rosenberg RN, White CL, 3rd, Ferrer I, Llado A, Neumann M, Kretzschmar HA, Hulette CM, Welsh-Bohmer KA, Miller BL, Alzualde A, Lopez de Munain A, McKee AC, Gearing M, Levey AI, Lah JJ, Hardy J, Rohrer JD, Lashley T, Mackenzie IR, Feldman HH, Hamilton RL, Dekosky ST, van der Zee J, Kumar-Singh S, Van Broeckhoven C, Mayeux R, Vonsattel JP, Troncoso JC, Kril JJ, Kwok JB, Halliday GM, Bird TD, Ince PG, Shaw PJ, Cairns NJ, Morris JC, McLean CA, DeCarli C, Ellis WG, Freeman SH, Frosch MP, Growdon JH, Perl DP, Sano M, Bennett DA, Schneider JA, Beach TG, Reiman EM, Woodruff BK, Cummings J, Vinters HV, Miller CA, Chui HC, Alafuzoff I, Hartikainen P, Seilhean D, Galasko D, Masliah E, Cotman CW, Tunon MT, Martinez MC, Munoz DG, Carroll SL, Marson D, Riederer PF, Bogdanovic N, Schellenberg GD, Hakonarson H, Trojanowski JQ, Lee VM (2010) Common variants at 7p21 are associated with frontotemporal lobar degeneration with TDP-43 inclusions. Nat Genet 42:234–239. doi:10.1038/ng.536

89. van der Zee J, Van Langenhove T, Kleinberger G, Sleegers K, Engelborghs S, Vandenberghe R, Santens P, Van den Broeck M, Joris G, Brys J, Mattheijssens M, Peeters K, Cras P, De Deyn PP, Cruts M, Van Broeckhoven C (2011) TMEM106B is associated with frontotemporal lobar degeneration in a clinically diagnosed patient cohort. Brain 134:808–815. doi:10.1093/brain/awr007

90. Vass R, Ashbridge E, Geser F, Hu WT, Grossman M, Clay-Falcone D, Elman L, McCluskey L, Lee VM, Van Deerlin VM, Trojanowski JQ, Chen-Plotkin AS (2011) Risk genotypes at TMEM106B are associated with cognitive impairment in amyotrophic lateral sclerosis. Acta Neuropathol 121:373–380. doi:10.1007/s00401-010-0782-y

91. Wagner OI, Ascano J, Tokito M, Leterrier JF, Janmey PA, Holzbaur EL (2004) The interaction of neurofilaments with the microtubule motor cytoplasmic dynein. Mol Biol Cell 15:5092–5100. doi:10.1091/mbc.e04-05-0401

92. Wang Y, Mandelkow E (2012) Degradation of tau protein by autophagy and proteasomal pathways. Biochem Soc Trans 40:644–652. doi:10.1042/BST20120071

93. Wes PD, Easton A, Corradi J, Barten DM, Devidze N, DeCarr LB, Truong A, He A, Barrezueta NX, Polson C, Bourin C, Flynn ME, Keenan S, Lidge R, Meredith J, Natale J, Sankaranarayanan S, Cadelina GW, Albright CF, Cacace AM (2014) Tau overexpression impacts a neuroinflammation gene expression network perturbed in Alzheimer’s disease. PLoS One 9:e106050. doi:10.1371/journal.pone.0106050

94. Xu Z, Cork LC, Griffin JW, Cleveland DW (1993) Increased expression of neurofilament subunit NF-L produces morphological alterations that resemble the pathology of human motor neuron disease. Cell 73:23–33. doi:10.1016/0092-8674(93)90157-l

95. Yan H, Kubisiak T, Ji H, Xiao J, Wang J, Burmeister M (2018) The recurrent mutation in TMEM106B also causes hypomyelinating leukodystrophy in China and is a CpG hot spot. Brain. doi:10.1093/brain/awy029

96. Yoshiyama Y, Higuchi M, Zhang B, Huang SM, Iwata N, Saido TC, Maeda J, Suhara T, Trojanowski JQ, Lee VM (2007) Synapse loss and microglial activation precede tangles in a P301S tauopathy mouse model. Neuron 53:337–351. doi:10.1016/j.neuron.2007.01.010

97. Yu L, De Jager PL, Yang J, Trojanowski JQ, Bennett DA, Schneider JA (2015) The TMEM106B locus and TDP-43 pathology in older persons without FTLD. Neurology 84:927–934. doi:10.1212/WNL.0000000000001313

98. Yuan A, Nixon RA (2021) Neurofilament Proteins as Biomarkers to Monitor Neurological Diseases and the Efficacy of Therapies. Front Neurosci 15:689938. doi:10.3389/fnins.2021.689938

99. Yuan A, Rao MV, Veeranna, Nixon RA (2017) Neurofilaments and Neurofilament Proteins in Health and Disease. Cold Spring Harb Perspect Biol 9. doi:10.1101/cshperspect.a018309

100. Zhang L, Sheng R, Qin Z (2009) The lysosome and neurodegenerative diseases. Acta Biochim Biophys Sin (Shanghai) 41:437–445. doi:10.1093/abbs/gmp031

101. Zhang T, Pang W, Feng T, Guo J, Wu K, Nunez Santos M, Arthanarisami A, Nana AL, Nguyen Q, Kim PJ, Jankowsky JL, Seeley WW, Hu F (2023) TMEM106B regulates microglial proliferation and survival in response to demyelination. Sci Adv 9:eadd2676. doi:10.1126/sciadv.add2676

102. Zhu B, Liu Y, Hwang S, Archuleta K, Huang H, Campos A, Murad R, Pina-Crespo J, Xu H, Huang TY (2022) Trem2 deletion enhances tau dispersion and pathology through microglia exosomes. Mol Neurodegener 17:58. doi:10.1186/s13024-022-00562-8

