## Supplementary material for "Loss of TMEM106B exacerbates Tau pathology and neurodegeneration in PS19 mice": Fig. S1-S8

**Supplemental Figures**

**Supplemental Figure 1. Loss of TMEM106B decreases the protein levels of NeuN, but increases the levels of cleaved caspase-3 in PS19 mice**

**Supplemental Figure 2. TMEM106B deficiency leads to the accumulation of sarkosyl-insoluble hTau and phosphorylated hTau, but does not affect the distribution of hTau in young PS19 mice**

**Supplemental Figure 3. Loss of TMEM106B in mice does not affect the level of endogenous mTau and phosphorylated mTau**

**Supplemental Figure 4. Deletion of TMEM106B leads to the accumulation of p-NF-H/M, NF-H and NF-L in neurons in PS19 mice**

**Supplemental Figure 5. TMEM106B deficiency results in increased GFAP protein levels.**

**Supplemental Figure 6. TMEM106B deficiency does not affect microglia activation in the young PS19 mice.**

**Supplemental Figure 7. TMEM106B ablation causes slight changes in lysosomal protein levels but does not affect the level of CathD in astrocytes in PS19 mice.**

**Supplemental Figure 8. Mutant hTau does not accumulate in *Tmem106b^-/-^* microglia and astrocytes.**

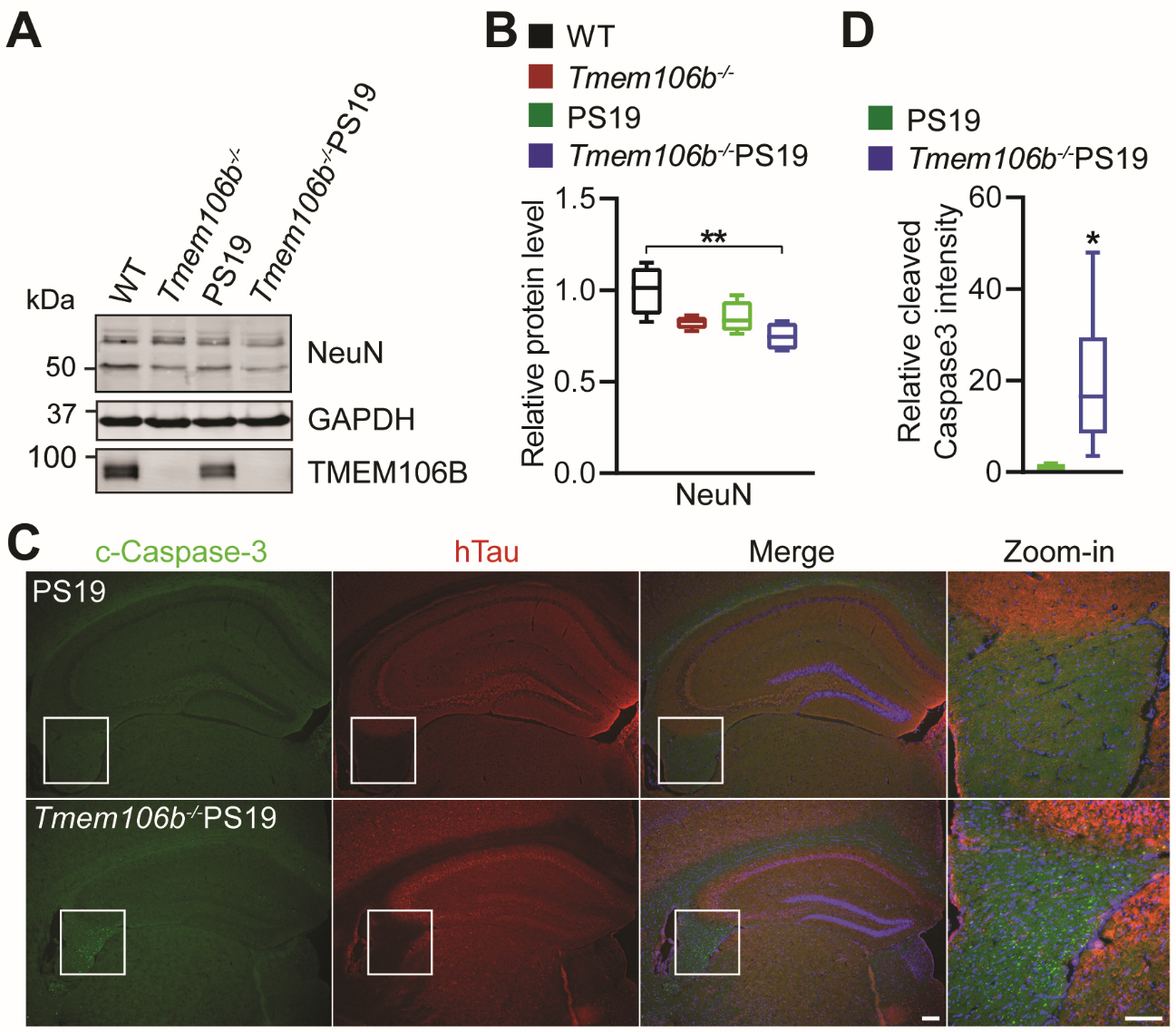

**Supplemental Figure 1. Loss of TMEM106B decreases the protein levels of NeuN, but increases the levels of cleaved caspase-3 in PS19 mice**

**(A, B)** Western blot analysis of NeuN, TMEM106B, and GAPDH in sarkosyl-soluble fraction extracted from the brain of 8.5-month-old WT, *Tmem106b^-/-^*, PS19, and *Tmem106b^-/-^* PS19 mice. Relative levels of indicated proteins were quantified in B. n=4. Data presented as mean ± SEM. One-way ANOVA tests with Bonferroni’s multiple comparisons: **, p<0.01. **(C, D)** Representative images of cleaved caspase-3 (c-Caspase-3) and hTau co-immunostaining in the hippocampus sections of 8.5-month-old PS19 and *Tmem106b^-/-^* PS19 mice. Zoom-in images show the fornix region in hippocampus. Relative cleaved caspase-3 intensity in hippocampus was quantified in D. n=6. Data presented as mean ± SEM. Unpaired Student’s t test: *, p<0.05. Scale bar: 100 µm.

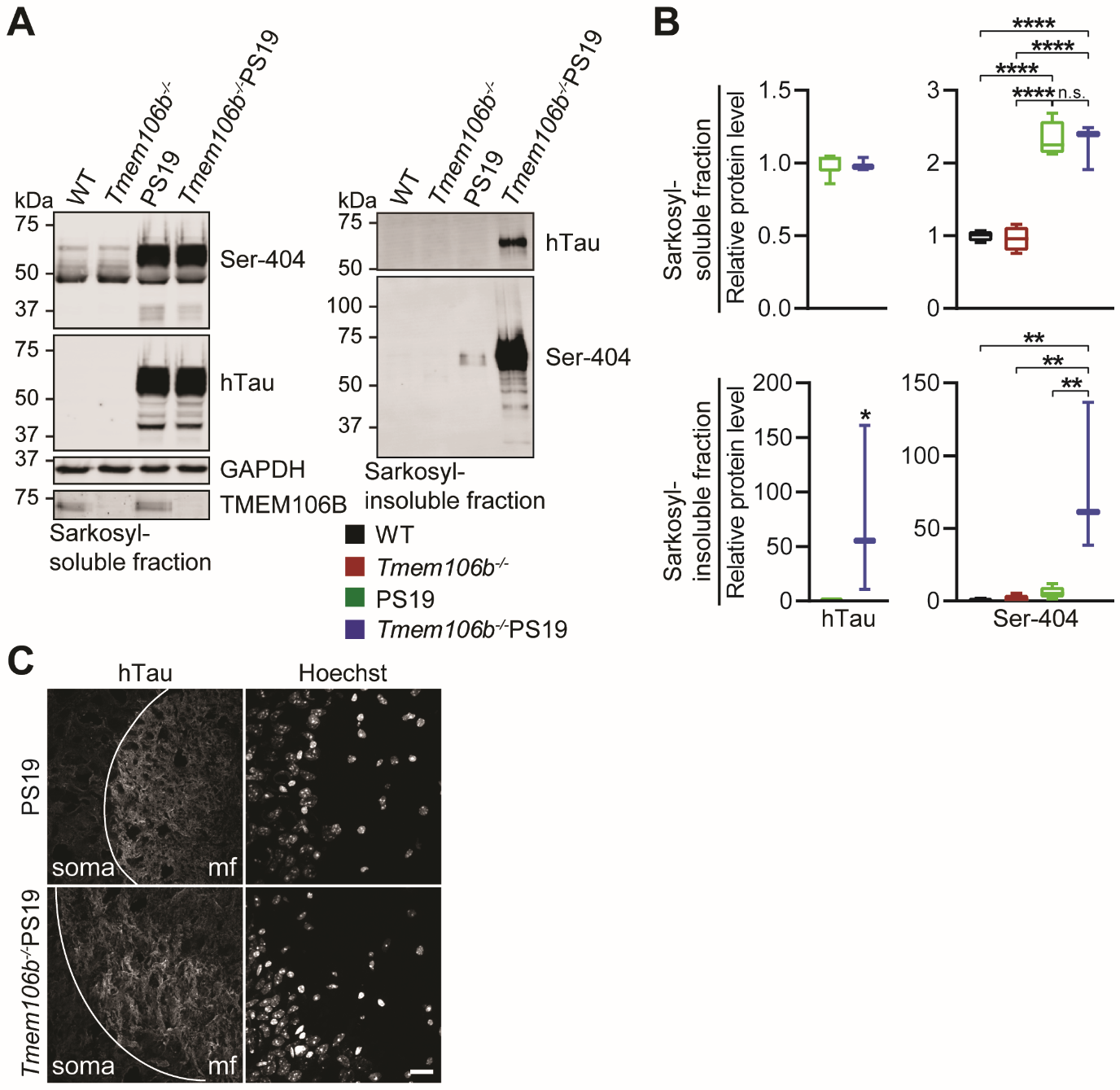

**Supplemental Figure 2. TMEM106B deficiency leads to the accumulation of sarkosyl-insoluble hTau and phosphorylated hTau, but does not affect the distribution of hTau in young PS19 mice**

**(A, B)** Western blot analysis of hTau, phosphorylated Tau (Ser-404), TMEM106B, and GAPDH in sarkosyl-soluble fraction and sarkosyl-insoluble fraction extracted from the hippocampus of 5 to 5.4-month-old WT, *Tmem106b^-/-^*, PS19, and *Tmem106b^-/-^* PS19 mice. Relative protein levels of indicated proteins were quantified in B. n=3-8. Data presented as mean ± SEM. One-way ANOVA tests with Bonferroni’s multiple comparisons: **, p<0.01; ****, p<0.0001. n.s.: non-significant. Unpaired Student’s t test for two groups comparison: *, p<0.05. **(C)** Representative confocal microscope images of hTau immunostaining in hippocampal CA3 region of 5 to 5.4-month-old PS19 and *Tmem106b^-/-^* PS19 mice brain sections. mf: mossy fibers. Scale bar: 20 µm.

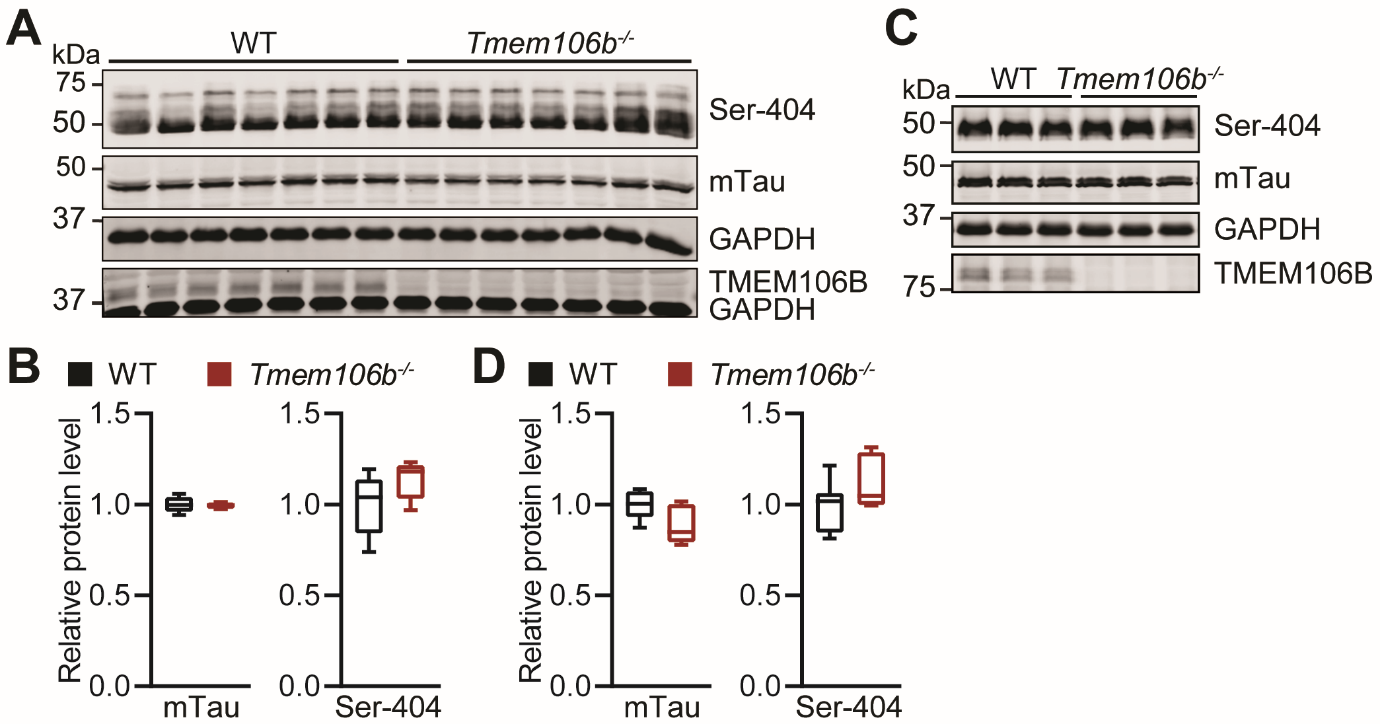

**Supplemental Figure 3. Loss of TMEM106B in mice does not affect the level of endogenous mTau and phosphorylated mTau.**

**(A, B)** Western blot analysis of mTau, phosphorylated mTau (Ser-404), TMEM106B, and GAPDH in RIPA-soluble fraction of hippocampus from 6-month-old WT and *Tmem106b^-/-^* mice. Relative levels of indicated proteins were quantified in B. n=7. Data presented as mean ± SEM. Unpaired Student’s t test for two groups comparison.

**(C, D)** Western blot analysis of mTau, phosphorylated Tau (Ser-404), TMEM106B, and GAPDH in RIPA-soluble fraction from 16-month-old WT and *Tmem106b^-/-^* mouse brain. Relative levels of indicated proteins were quantified in D. n=5. Data presented as mean ± SEM. Unpaired Student’s t test for two groups comparison.

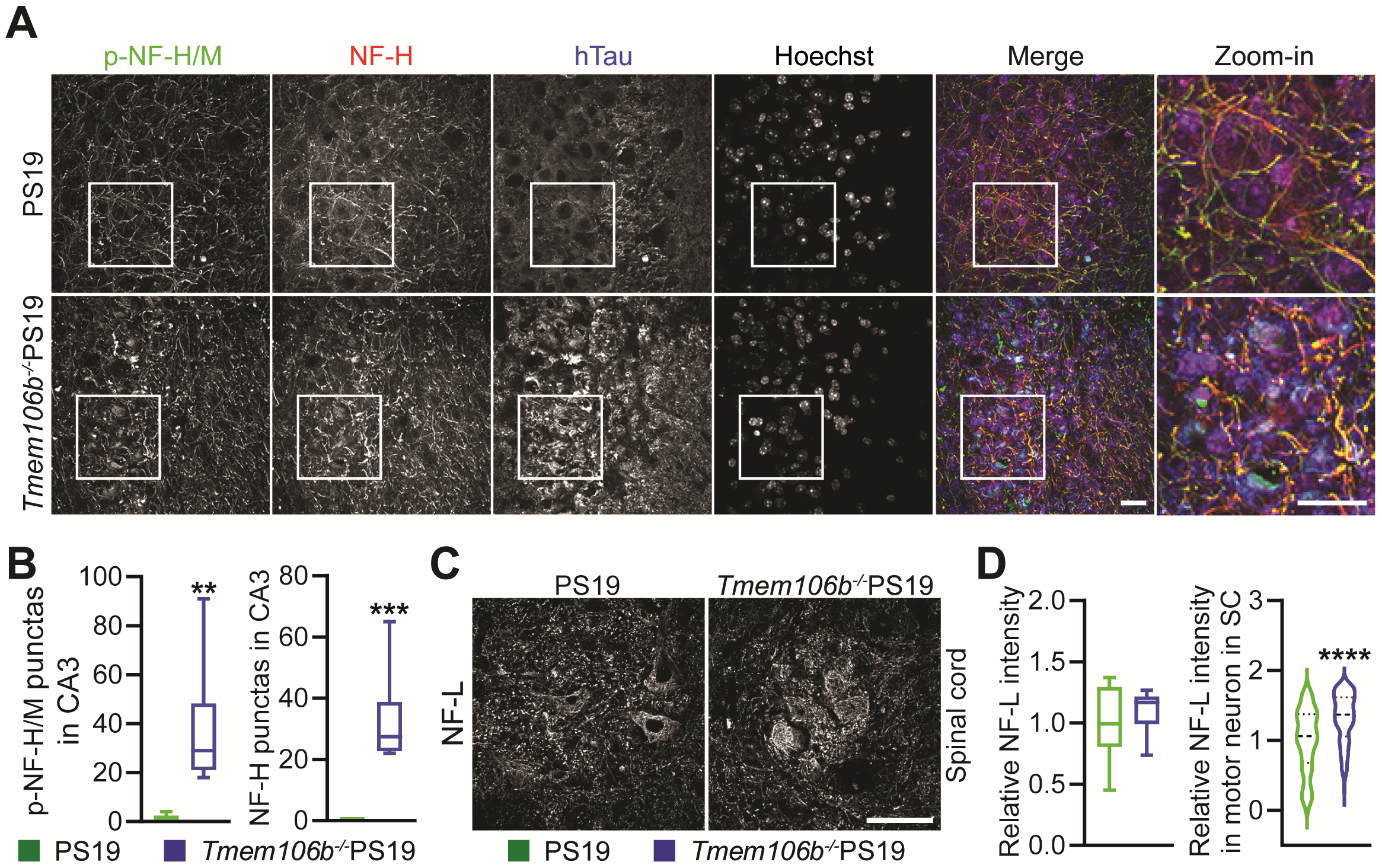

**Supplemental Figure 4. Deletion of TMEM106B leads to the accumulation of p-NF-H/M, NF-H and NF-L in neurons in PS19 mice**

**(A, B)** Representative image of p-NF-H/M, NF-H and hTau co-immunostaining in hippocampal CA3 region of 8.5-month-old PS19 and *Tmem106b^-/-^* PS19 mice. Relative number of fluorescent punctas of p-NF-H/M or NF-H was quantified in B. n=6. Data presented as mean ± SEM. Unpaired Student’s t test: **, p<0.01; ***, p<0.001. Scale bar: 20 µm.

**(C, D)** Representative image of NF-L immunostaining in the spinal cord of 8.5-month-old PS19 and *Tmem106b^-/-^* PS19 mice. Total NF-L intensity and relative fluorescence intensity of NF-L in motor neurons were quantified in D. n=6; Number of motor neuron =134-159. Data presented as mean ± SEM. Unpaired Student’s t test: ****, p<0.0001. n.s.: non-significant. Scale bar: 20 µm.

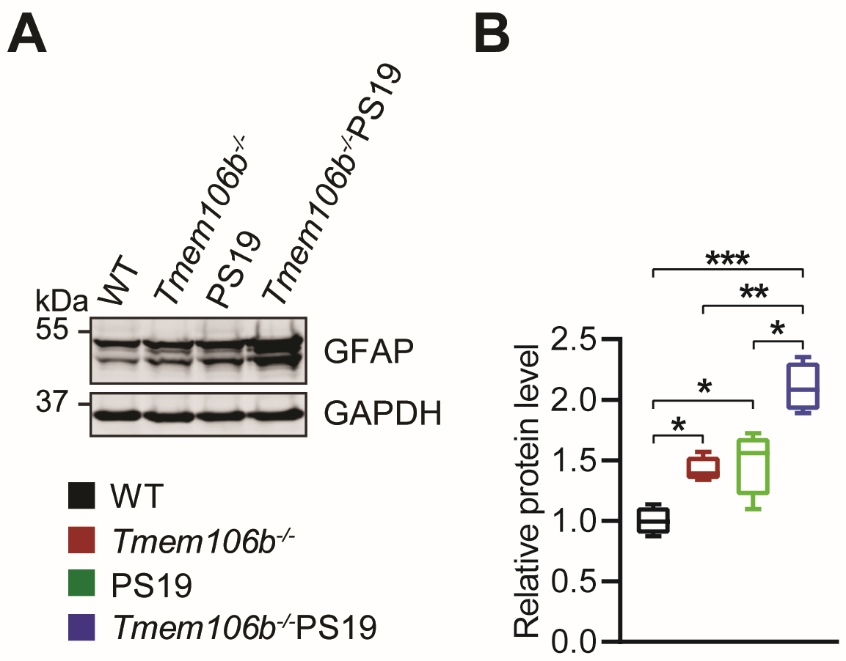

**Supplemental Figure 5. TMEM106B deficiency results in increased GFAP protein levels.**

**(A, B)** Western blot analysis of GFAP and GAPDH in sarkosyl-soluble fraction extracted from the brain of 8.5-month-old WT, *Tmem106b^-/-^*, PS19, and *Tmem106b^-/-^* PS19 mice. Relative GFAP protein level was quantified in B. n=4. Data presented as mean ± SEM. One-way ANOVA tests with Bonferroni’s multiple comparisons: *, p<0.05; **, p<0.01; ***, p<0.001.

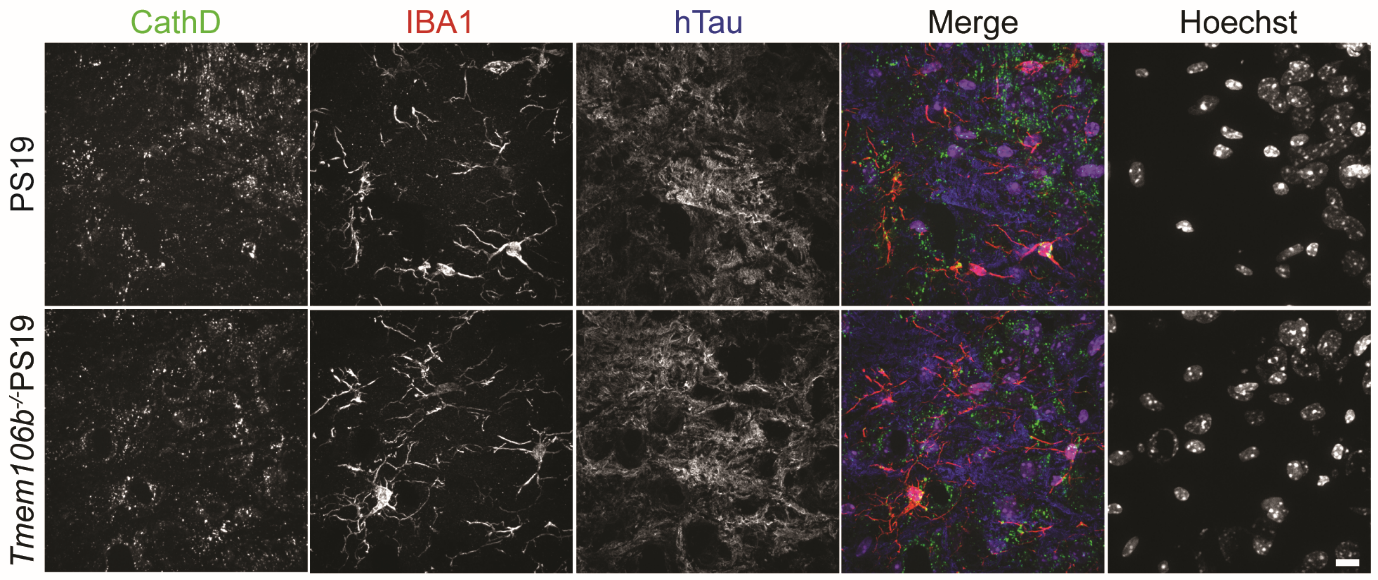

**Supplemental Figure 6. TMEM106B deficiency does not affect microglia activation in the young PS19 mice.**

Representative images of IBA1, CathD and hTau co-immunostaining in the hippocampus of 5-month-old PS19 and *Tmem106b^-/-^* PS19 mice. Scale bar: 10 µm.

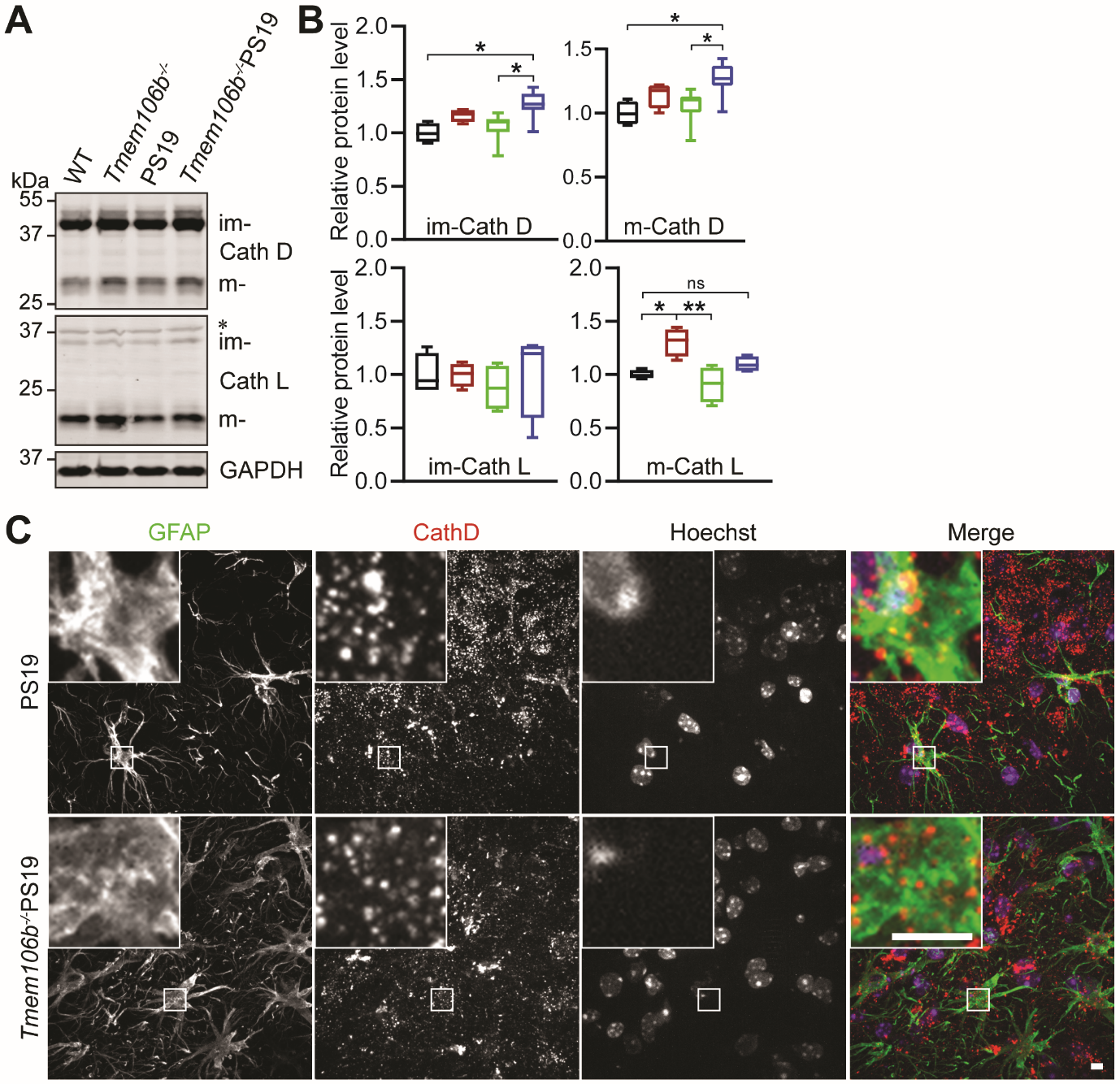

**Supplemental Figure 7. TMEM106B ablation causes slight changes in lysosomal protein levels but does not affect the level of CathD in astrocytes in PS19 mice.**

**(A, B)** Western blot analysis of lysosomal proteins in sarkosyl-soluble fraction extracted from the brain of 8.5-month-old WT, *Tmem106b^-/-^*, PS19, and *Tmem106b^-/-^* PS19 mice. Asterisk indicates non-specific bands. Relative levels of indicated proteins were quantified in B. n=4-6. Data presented as mean ± SEM. One-way ANOVA tests with Bonferroni’s multiple comparisons: *, p<0.05; **, p<0.01; n.s., non-significant. (**C**) Representative images of GFAP and CathD co-immunostaining in hippocampal CA3 region of 8.5-month-old PS19 and *Tmem106b^-/-^* PS19 mice. Scale bar: 5 µm.

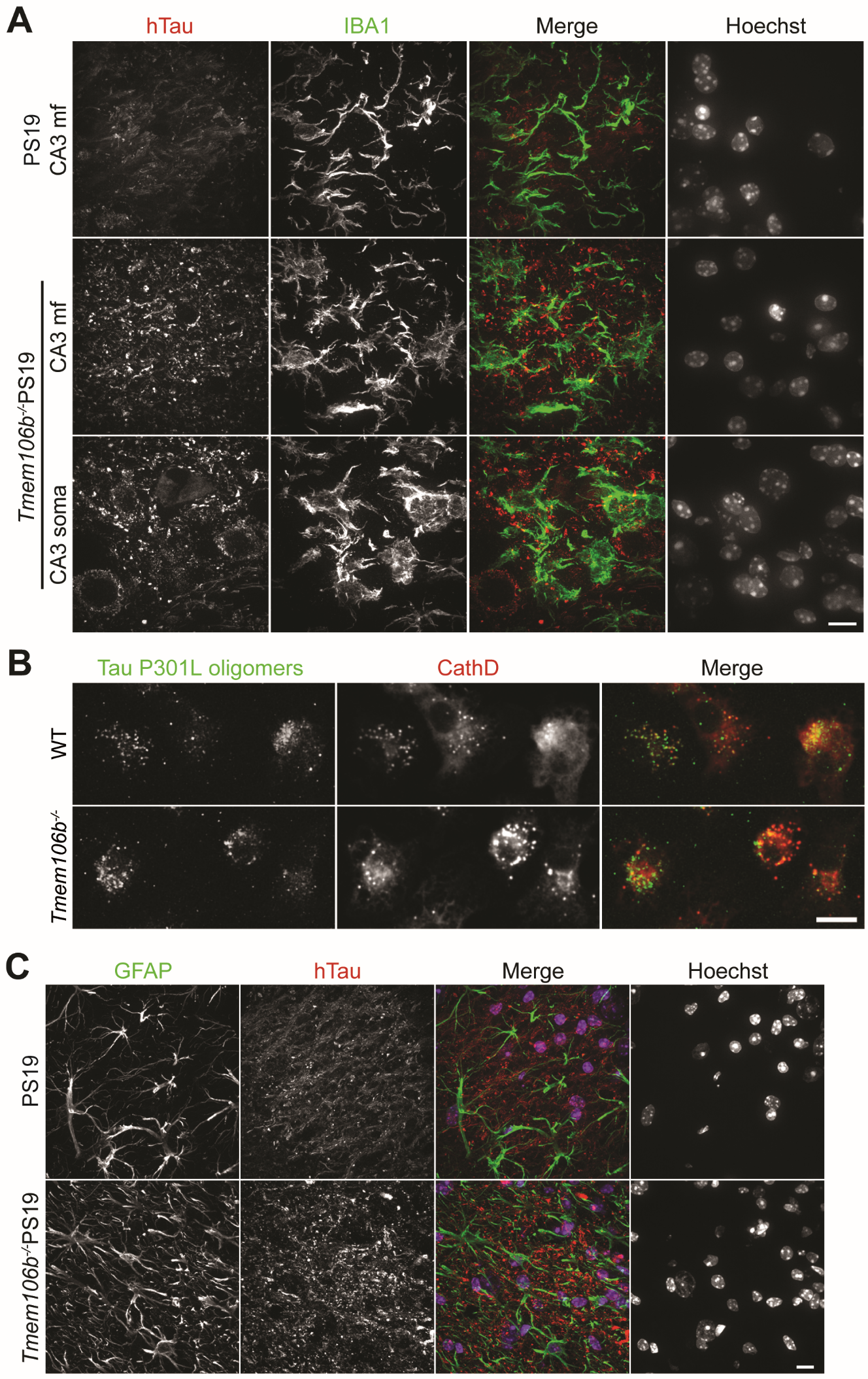

**Supplemental Figure 8. Mutant hTau does not accumulate in *Tmem106b^-/-^* microglia and astrocytes.**

**(A)** Representative image of hTau and IBA1 co-immunostaining in hippocampal CA3 region of 8.5-month-old PS19 and *Tmem106b^-/-^* PS19 mice. Scale bar: 10 µm.

**(B)** Representative image of CathD and IBA1 co-immunostaining in primary cultured microglia treated with 100 nM DyLight 488-Tau P301L oligomer. Scale bar: 10 µm.

**(C)** Representative image of hTau and GFAP co-immunostaining in hippocampal CA3 region of 8.5-month-old PS19 and *Tmem106b^-/-^* PS19 mice. Scale bar: 10 µm.
